# CRISPR/Cas9 screen reveals a role of purine synthesis for estrogen receptor α activity and tamoxifen resistance of breast cancer cells

**DOI:** 10.1101/2022.06.03.494664

**Authors:** Dina Hany, Vasiliki Vafeiadou, Didier Picard

## Abstract

In breast cancer, resistance to endocrine therapies that target estrogen receptor α (ERα), such as tamoxifen and fulvestrant, remains a major clinical problem. Whether and how ERα+ breast cancers switch from being estrogen-dependent to -independent remains unclear. With a genome-wide CRISPR/Cas9 knockout screen, we identified new biomarkers and potential therapeutic targets of endocrine resistance. We demonstrate that high levels of PAICS, an enzyme involved in the *de novo* biosynthesis of purines, can shift the balance of ERα activity to be more estrogen-independent and tamoxifen-resistant. We indicate that this is due to an elevated activity of cAMP-activated protein kinase A and mammalian target of rapamycin, kinases known to phosphorylate ERα specifically and to stimulate its activity. Genetic or pharmacological targeting of PAICS sensitizes tamoxifen-resistant cells to tamoxifen. Based on these findings, we propose the combined targeting of PAICS and ERα as a new, effective, and potentially safe therapeutic regimen.

## Introduction

Breast cancer remains a major threat for women worldwide. According to the latest update of global cancer statistics, female breast cancer presented the highest incidence rate with an estimated 2.3 million new cases (11.7%) in 2020, unprecedentedly surpassing lung cancer (11.4%). Among women, breast cancer has the highest 5-year prevalence (30.3%) and the highest mortality rate (15.5%) compared to other types of cancer (*1*). Breast cancer is globally classified based on clinical features and molecular biomarkers. Invasive ductal carcinoma represents the most common histologic subtype of breast cancer (*2*). Molecular classification is based on the expression of the steroid receptors estrogen receptor α (ERα) and progesterone receptor (PR), the human epidermal growth factor receptor 2 (HER2), and the proliferation marker Ki-67. Luminal breast cancer that expresses ERα represents approximately 70% and includes the subtypes luminal A (ERα+, PR+, HER2-, and low Ki-67) and luminal B (ERα+, PR+/−, HER2−/+, and high Ki-67) (*2, 3*).

ERα plays a central role in promoting survival, proliferation, invasion, and metastasis of breast cancer. ERα is a ligand-regulated transcription factor, which contains two activation functions, an N-terminal ligand-independent activation function 1 (AF-1) and a ligand-dependent activation function 2 (AF-2), which is associated with the C-terminal hormone binding domain (*4*). AF-1 and AF-2 can mediate transcriptional activation of ERα both independently and synergistically (*4*). Upon estrogen binding, ERα dissociates from molecular chaperones, dimerizes, and binds with coregulators to specific palindromic DNA sequences called estrogen response elements (EREs) leading to the induction or repression of target genes (*5, 6*). Ligand-bound ERα can also be tethered indirectly to the response elements of other transcription factors, such as the activator protein 1 (AP-1), nuclear factor kappa B (NFκB), and stimulating protein 1, modulating their activities (*7*). In a non-genomic manner, the binding of estrogens to a membrane-associated subset of ERα molecules can lead to the rapid activation of various protein kinase signaling pathways, such as those involving phosphatidylinositol 3-kinase (PI3K), protein kinase B (hereafter referred to as Akt), mammalian target of rapamycin (mTOR), and mitogen-activated protein kinases (MAPKs), causing both cytoplasmic and genomic responses (*8*). ERα can also be activated as a transcription factor in the absence of estrogens by a wide variety of signal transduction pathways (see updated list at https://www.picard.ch/ downloads/Factors.pdf), including the cyclic adenosine monophosphate (cAMP)/protein kinase A (PKA) and the PI3K/Akt/mTOR pathways (*9, 10*). Phosphorylation of ERα occurs at multiple sites and regulates ERα functions such as sensitivity to hormones (*11*), DNA binding (*12, 13*), and regulation of target gene transcription (*14*). It plays an important role in estrogen-dependent and -independent activation of ERα and in the response to endocrine therapy of breast cancer (*4, 9, 10, 14–16*). For example, S305 of ERα has been shown to be the main phosphorylation target for cAMP-activated PKA, and it is associated with an enhanced agonistic behavior of tamoxifen and resistance to treatment with tamoxifen (*10, 15–17*).

For decades, targeting ERα has represented the gold standard strategy for therapy of ERα+ breast cancer. Therapeutic targeting of ERα includes inhibition of the synthesis of estrogens using aromatase inhibitors, such as anastrozole and exemestane, competing with estrogen binding using the pure anti-estrogen fulvestrant/ICI182,780 (ICI) or the partial agonist/antagonist tamoxifen. The former prevents the activation of both AF-1 and AF-2, whereas the latter inhibits the activation of AF-2 but not AF-1 (*4*). Tamoxifen represents the backbone of adjuvant endocrine therapy for both pre- and post-menopausal women with breast cancer (*18*). It has proven to be relatively well-tolerated and highly efficient, even though drug resistance eventually develops. As tamoxifen resistance represents a significant problem in clinical practice, it remains a concern for basic and translational research. Many mechanisms have been described as being responsible for the escape of cancer cells from the cytotoxic effects of tamoxifen, including loss or deregulation of ERα expression and alterations of signaling pathways responsible for survival, proliferation, apoptosis, and stress responses (*9, 10, 15, 16, 19–21*). Metabolic rewiring plays a crucial role in cellular growth, survival, and proliferation to generate an adequate amount of adenosine triphosphate (ATP) and to fuel the biosynthesis of macromolecules in breast cancer (*22*). There is evidence supporting the role of reprogramming lipid, glucose (*23, 24*), and amino acid (*25, 26*) metabolism in the resistance to endocrine therapy. As an example, metabolic upregulation in the amino acids serine and aspartate/glutamate are associated with increased nucleotide biosynthesis, and resistance to tamoxifen (*25, 26*).

Despite a substantial number of studies, failure of endocrine therapy remains a major clinical challenge. Intratumor heterogeneity (*27*) and interpatient genetic and epigenetic tumor variabilities (*28*) might contribute to the diverse responses to drugs in breast cancer (*20*). A switch of primary ERα+ breast cancer from being ligand-dependent to ligand-independent represents a key cause that remains mechanistically obscure. Therefore, the identification of the relevant genetic factors or biomarkers can help improve the understanding of the underlying mechanisms and hence, orienting the therapeutic choices. For this study, our aim was to identify new factors that play a role in ligand-dependent and -independent signaling of ERα, and in the responses of breast cancer to endocrine therapy with tamoxifen or ICI. Genetic screens using the clustered, regularly interspaced, short palindromic repeats (CRISPR) system using the RNA-guided endonuclease Cas9 represent powerful tools for the discovery of essential genes, drug targets, or biomarkers of drug resistance (*29–32*). Screens with the tamoxifen-sensitive breast cancer cell lines MCF7 and T47D identified gene deletions, such as that of *CSK*, mediating estrogen-independent growth (*32*). The expression of *ARID1A* and components of the SWI/SNF complex were identified as prerequisites for the response of MCF7 cells to 4-hydroxytamoxifen (4-OHT), the major active metabolite of tamoxifen, and to ICI (*30*). A chemogenetic screen using an inhibitor of the ERα coactivator SRC-3 in MCF7 cells identified several new potential drug targets (*29*). Moreover, a specific CRISPR/Cas9 screen targeting CTCF binding sites in the vicinity of ERα-bound enhancers identified ERα target genes such as *PREX1* as being vital for ERα function (*33*). Despite all these previous efforts, a more comprehensive identification and characterization of the genetic determinants of ERα-stimulated proliferation of breast cancer cells remains to be done.

We therefore performed parallel genome-wide CRISPR/Cas9 knockout screens in the absence or presence of estrogen, upon estrogen-independent activation of ERα by cAMP-activated PKA, and in the presence of the ERα antagonists 4-OHT and ICI. Importantly, we used a MCF7 variant (MCF7-V) for our screens, which is relatively more 4-OHT-tolerant and displays more robust ERα responses (*34*). Our results contribute to a more complete understanding of the genes responsible for estrogen-independent and antiestrogen-resistant growth of breast cancer. Here we highlight the role of *PAICS*, one of the top hits in our screens. It encodes a bifunctional enzyme, which is involved in the *de novo* biosynthesis of purine nucleotides and carries out two successive reactions as a phosphoribosylaminoimidazole carboxylase and as a phosphoribosylamino-imidazole-succinocarboxamide synthetase (*35*). Our work connects the biosynthesis and availability of purines to endocrine resistance in breast cancer.

## Results

### CRISPR/Cas9 screens identify determinants of 4-OHT and fulvestrant responses in ERα+ breast cancer cells

We used a genome-wide CRSIPR/Cas9 knockout screen approach in which, 17β-estradiol (E2) and dibutyryl-cAMP (db-cAMP), a cell-permeable version of cAMP, were used as inducers of cell proliferation, whereas 4-OHT and ICI were used as inhibitors of cell proliferation of MCF7-V cells (fig. S1, A and B). We used the human Brunello CRISPR knockout pooled library, which targets 19,114 functional genes of the human genome with 3 or 4 sgRNAs each (*36*). The library was amplified and validated for its complexity by next-generation sequencing (NGS) (fig. S1, C and D). A pool of viral particles was prepared to infect MCF7-V cells (Fig. 1A and fig. S1D). Transduced cells were then selected with puromycin, pooled, and subjected to the following conditions: only vehicle (for the negative control), E2, db-cAMP, E2 + 4-OHT, E2 + ICI, E2 + 4-OHT + db-cAMP, and E2 + ICI + db-cAMP (Fig. 1A and fig. S1, B and D). A low concentration of E2 of 100 pM was used to induce ERα-mediated cell proliferation, whereas ERα antagonists were used at 100 nM, a dose that is sufficiently high to counteract E2 (fig. S1, A and B), and yet relatively low to avoid unspecific effects. Before treatments, one sample of the pooled cells was collected to represent the control condition (Control “T0”) (Fig. 1A and fig. S1D). Note that our choice of treatments allows both positive and negative selections. For example, in the case of E2 and db-cAMP, which both enhance the growth and proliferation of ERα+ MCF7-V cells (fig. S1, A and B), a gene whose knockout impairs proliferation, and hence is negatively selected, can be considered as activator of the E2 or db-cAMP growth stimulatory pathways. In contrast, in the case of growth inhibitors like 4-OHT and ICI, a gene whose knockout causes the cells to survive the inhibitory effects of the drugs (i.e. is positively selected) might be responsible for drug sensitivity. By NGS, we identified the enrichment and depletion of sgRNAs for each treatment condition. Normalized counts of sgRNAs and the calculated β-scores are available in separate files (data S1 and data S2, respectively). There are approximately 600 genes whose sgRNA sequences were strongly depleted regardless of the treatment. We removed them “essential genes” (data S3) from further analyses. Next, a set of 58 genes was identified and hierarchically clustered into 15 distinct groups (Fig. 1B and data S4).

**Fig. 1.**
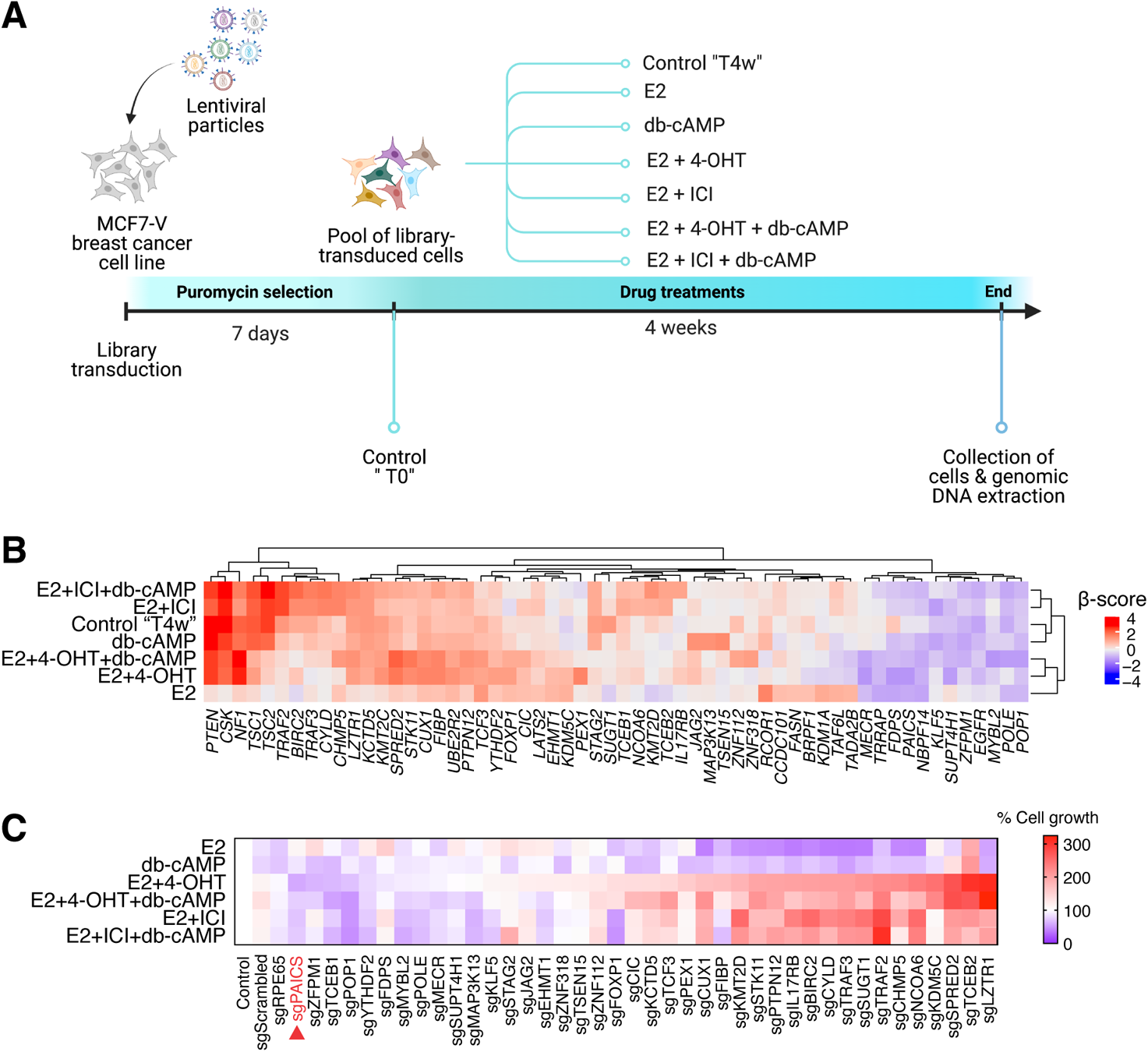
CRISPR/Cas9 screen identifies determinants of 4-OHT and ICI responses in ERα+ breast cancer cells. **(A)** Schematic representation and timeline of the primary CRISPR/Cas9 screen. Cells collected before drug treatments at time 0 (Control “T0”) and vehicle-treated cells collected after 4 weeks (Control “T4w”) were used as controls. **(B)** Heatmap representation of the hierarchical cluster analysis done for the top 58 genes identified by the primary CRISPR/Cas9 screen. Values are based on the calculated β-score of the pooled sgRNA reads from 2 biological replicates. Red indicates enrichment and blue indicates depletion of the sgRNAs of the corresponding genes. Cut-off value: β-score ≤ - 0.3 or ≥ 0.3, FDR ≤ 0.1 or ≥ −0.1. **(C)** Heatmap representation of % cell growth of individual knockouts of 40 genes selected for a secondary screen. Cell growth was measured by a crystal violet assay. Absorbances from each knockout cell line were first normalized to vehicle-treated control, and then to the corresponding values of non-transduced WT MCF7-V cells (set to 100%). Cells transduced with sgScrambled (expressing a scrambled sequence) or sgRPE65 (expressing a sgRNA targeting an unexpressed gene) represent two additional control cell lines. Values represent averages of 3 independent experiments. Red and blue indicate increased and reduced cell growth, respectively. Values were ordered ascendingly based on % cell growth with 4-OHT. Red arrow indicates sgPAICS as the knockout cell line with the highest sensitivity to 4-OHT.

They represent the top hits that show significant enrichment or depletion of sgRNA sequences in each condition as compared to the vehicle-treated control. Some of the top hits identified by our screen, including *PTEN*, *CSK*, *NF1*, and *TSC1/2* had previously been identified by a CRISPR/Cas9 screen that was performed with the ERα+ breast cancer cell lines MCF7 and T47D (*32*). Based on literature curation, a subset of 40 genes with potentially novel functions in the context of ERα+ breast cancer were identified and selected for further validation through a secondary screen. For each gene, we selected the sequence of one sgRNA present in the pooled Brunello library to produce a single knockout of the corresponding gene in MCF7-V cells. A polyclonal pool of knockout cells for each gene was subjected to the same treatments as in the primary screen and the results were represented as a heat map of cell growth (Fig. 1C). This analysis confirmed that out of the 40 selected genes, 33 might play a role in ERα signaling and/or 4-OHT or ICI responses. For example, loss of function of *LZTR1*, *TCEB2*, and *SPRED2* caused the cells to be significantly more resistant to 4-OHT and to a lesser extent to ICI (Fig. 1, B and C). We have recently shown that *SPRED2* deficiency is an important indicator of tamoxifen resistance because of ERα hyperactivation through the MAPK signaling pathway (*37*).

Moreover, the loss of functions of *BIRC2*, *TRAF2*, *TRAF3*, *CYLD*, and *CHMP5*, which all encode factors involved in TNFα-mediated activation of NFκB, were associated with significant resistance to ICI and to a lesser extent to 4-OHT (Fig. 1, B and C). In contrast, the loss of function of another set of genes, such as *PAICS*, *ZFPM1*, and *POP1*, enhanced the sensitivity to 4-OHT/ICI treatments (Fig. 1, B and C). We decided to focus on PAICS for the following.

### PAICS is upregulated in cancer and correlates with progression and poor clinical outcomes of ERα+ breast tumors

PAICS, a dual enzyme involved in the *de novo* biosynthesis of purines, appeared as the top hit of knockouts that sensitize MCF7-V cells to 4-OHT and ICI. First, to determine whether PAICS protein levels correlate with oncogenic characteristics, we compared them across multiple breast cancer cell lines (MCF7, MCF7-V, T47D, H3396, BT474, SKBR3, and MDA-MB-231) and the non-cancerous breast epithelial cell line MCF10A (fig. S2A). In addition, we compared several isogenic pairs of oncogenically transformed non-breast cell lines and their non-cancer parent derivatives. These included the non-transformed hTERT-immortalized retinal pigment epithelial cell line RPE1 along with its counterpart transformed by the H-Ras mutant G12V (RPE1 Ras) (fig. S2B), the non-tumorigenic lung epithelial cell line BEAS-2B and its v-Ha-Ras-transformed derivative BZR (fig. S2C), and human skin fibroblasts that were sequentially transformed by the stable overexpression of human telomerase (hTERT), hTERT + SV40 large T-antigen (hTERT LT), and hTERT + SV40 large T-antigen + H-RasG12V (hTERT LT Ras) (fig. S2D) (*38, 39*). We consistently observed higher PAICS protein levels in cancer cell lines as compared to their respective non-cancerous counterparts (fig. S2, A to D). The specific correlation between PAICS expression and breast cancer was further investigated using a human breast tissue microarray (TMA). Fluorescent immunostaining of the PAICS protein showed elevated expression in malignant breast tissues including intraductal and invasive ductal carcinomas as compared to non-malignant tissue biopsies including adjacent normal breast tissues and fibroadenomas, in both ERα+ and ERα-samples (Fig. 2, A to C). PAICS expression showed a strong correlation with the tumor progression based on the pathological diagnosis (Fig. 2, A and B), the expression of the cell proliferation marker Ki-67 (Fig. 2, D and E, and fig. S2, E and F), the tumor grade (Fig. 2F and fig. S2, G and H), and the “tumor, node, and metastasis” (TNM) stage (Fig. 2G, and fig. S2, I and J).

**Fig. 2.**
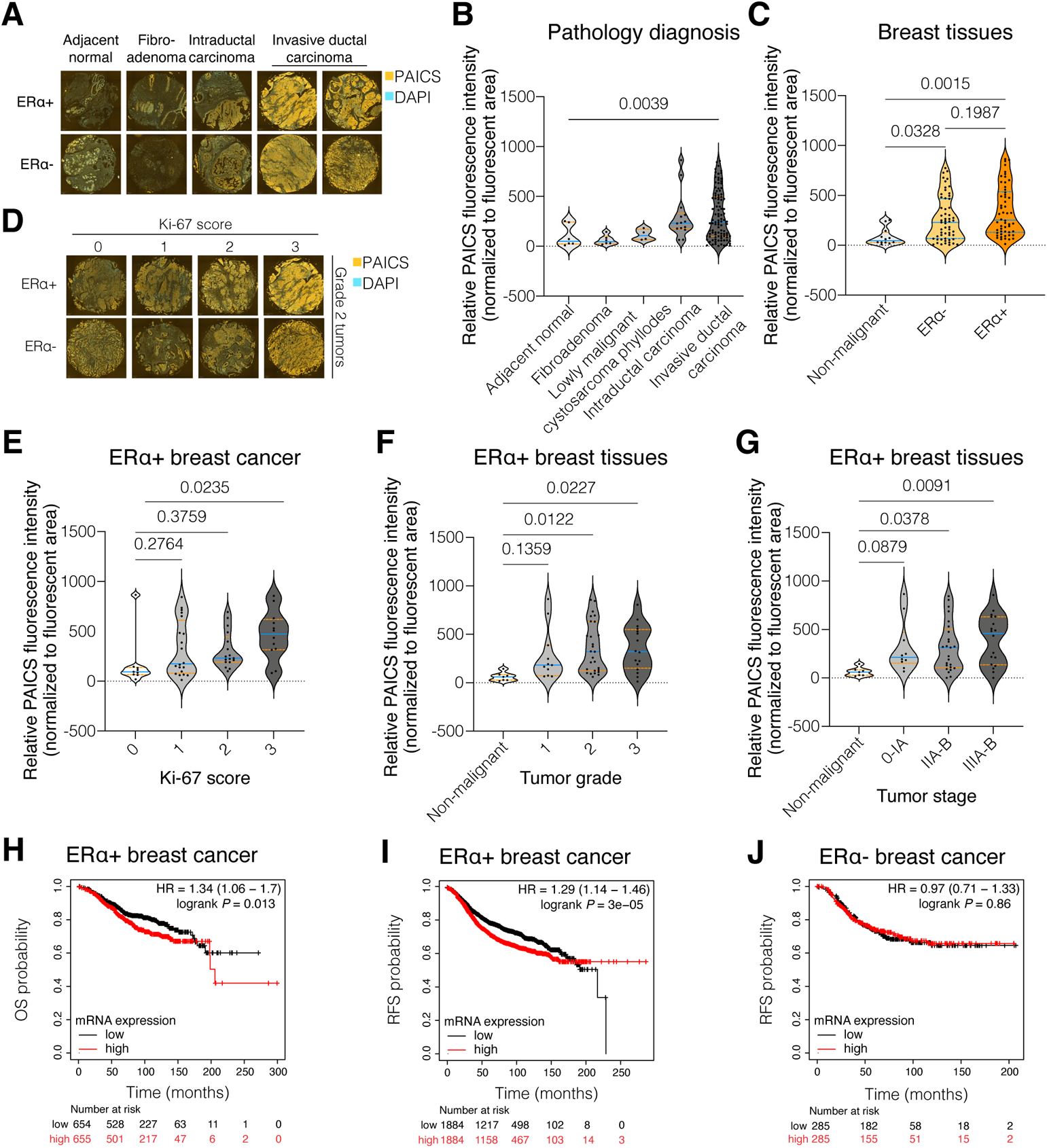
PAICS is upregulated in cancer and correlates with progression and poor clinical outcomes of ERα+ breast tumors. **(A and D)** Representative images from the TMA analysis of human breast tissue cores immunostained for PAICS (orange) and counterstained for the nuclei with DAPI (blue). Images representative of the pathological diagnosis (A) and Ki-67 score (D) among ERα+ and ERα-breast samples. **(B, C, and, E to G)** Violin plots of the means of the relative fluorescence intensities of PAICS per fluorescent unit area and its 95% confidence interval between cores, classified based on: pathology diagnosis (B), malignancy and ERα status (C), Ki-67 score of ERα+ breast cancer (E), tumor grade in ERα+ breast tissues (F), and TNM stage in ERα+ breast samples (G). The statistical significance between the groups was analyzed by one-way ANOVA, and *p*-values < 0.05 were considered statistically significant. **(H to J)** Kaplan-Meier plots for OS (H) and RFS (I) of ERα+ breast cancer, and OS (J) in ERα-ones, classified as tumors expressing high levels (red line) and low levels (black line) of *PAICS* mRNA.

More specifically, statistically significant correlations were associated with ERα+ (Fig. 2, E to G) rather than ERα-tumors (fig. S2, F, H, and J). Next, we used the Kaplan-Meier plotter (*40*) to search for the correlation between *PAICS* mRNA expression levels and survival of breast cancer patients. We found that *PAICS* gene expression is significantly correlated with poor overall survival (OS) and relapse-free survival (RFS) of patients with ERα+ breast tumors (Fig. 2, H and I). High *PAICS* mRNA levels are also correlated with poor outcomes in those patients who received tamoxifen therapy (fig. S2, K and L). In contrast, *PAICS* gene expression is not correlated with OS or RFS of ERα-breast cancer patients (Fig. 2J and fig. S2M). These results suggest that PAICS is highly expressed in cancer, can be a marker of breast tumor progression, and can determine disease outcomes in patients with ERα+ breast cancer.

### PAICS expression correlates with cell migration, estrogen-independent and tamoxifen-resistant proliferation of ERα+ breast cancer cells

To explore whether *PAICS* can function as an oncogene, we stably overexpressed a PAICS-EGFP fusion protein in the non-cancerous RPE1 cell line (fig. S3A). RPE1 cells overexpressing PAICS display more irregular, spreading, and protrusive shapes than the parent cells (fig. S3B). In a wound-healing assay, PAICS overexpression significantly enhanced cell migration and fastened healing of the wounded area as compared to WT non-transfected RPE1 (fig. S3, C and D). For further characterization with ERα+ breast cancer cells, we generated two individual knockdown clones of PAICS in MCF7-V cells using a CRISPR/Cas9 sgRNA lentiviral construct targeting *PAICS* (sgPAICS) (Fig. 3A). In parallel, cells were transduced with lentiviral particles for the expression of a sgRNA targeting the unexpressed gene *RPE65* as a negative control (sgNT). Since we could not identify a full knockout clone of PAICS, we hypothesize that some low level of PAICS may be essential for survival, similarly to the other members of the *de novo* purine biosynthesis pathway, such as GART (data S3). Two MCF7-V clones with stable overexpression of PAICS-EGFP were generated using the expression plasmid pPAICS-EGFP. These cellular clones were referred to as PAICS OE 45 and PAICS OE 25 (Fig. 3B). Another clone that was obtained in parallel but was not showing overexpression of PAICS by immunoblotting was used as a negative control (PAICS NC). Similarly, two PAICS-overexpressing clones were obtained in T47D cells and are referred to as PAICS OE 2 and PAICS OE 8 (fig. S3E). Reminiscent of the results obtained with PAICS-overexpressing RPE1 cells, MCF7-V cells showed enhanced cell migration when PAICS was overexpressed (Fig. 3, C and D, and fig. S3F), whereas no significant difference to WT cells was observed in the case of the knockdown of PAICS (fig. S3, G and H). Next, we determined more specifically whether the differential expression of PAICS affects the proliferation rate of ERα+ breast cancer cells. We found that its overexpression increases the proliferation of MCF7-V and T47D cells in the absence of E2 as well as in the presence of 4-OHT, but not ICI, by comparison with PAICS NC cells (Fig. 3, E to G, and fig. S3, I to K). PAICS deficiency caused MCF7-V cells to become more dependent on E2 for their proliferation (Fig. 3, H to J). Taken together, these results indicate that increased PAICS expression may stimulate the proliferation and migration of ERα+ breast cancer cells in the absence of estrogen, whereas PAICS deficiency restricts cell proliferation to the presence of estrogen.

**Fig. 3.**
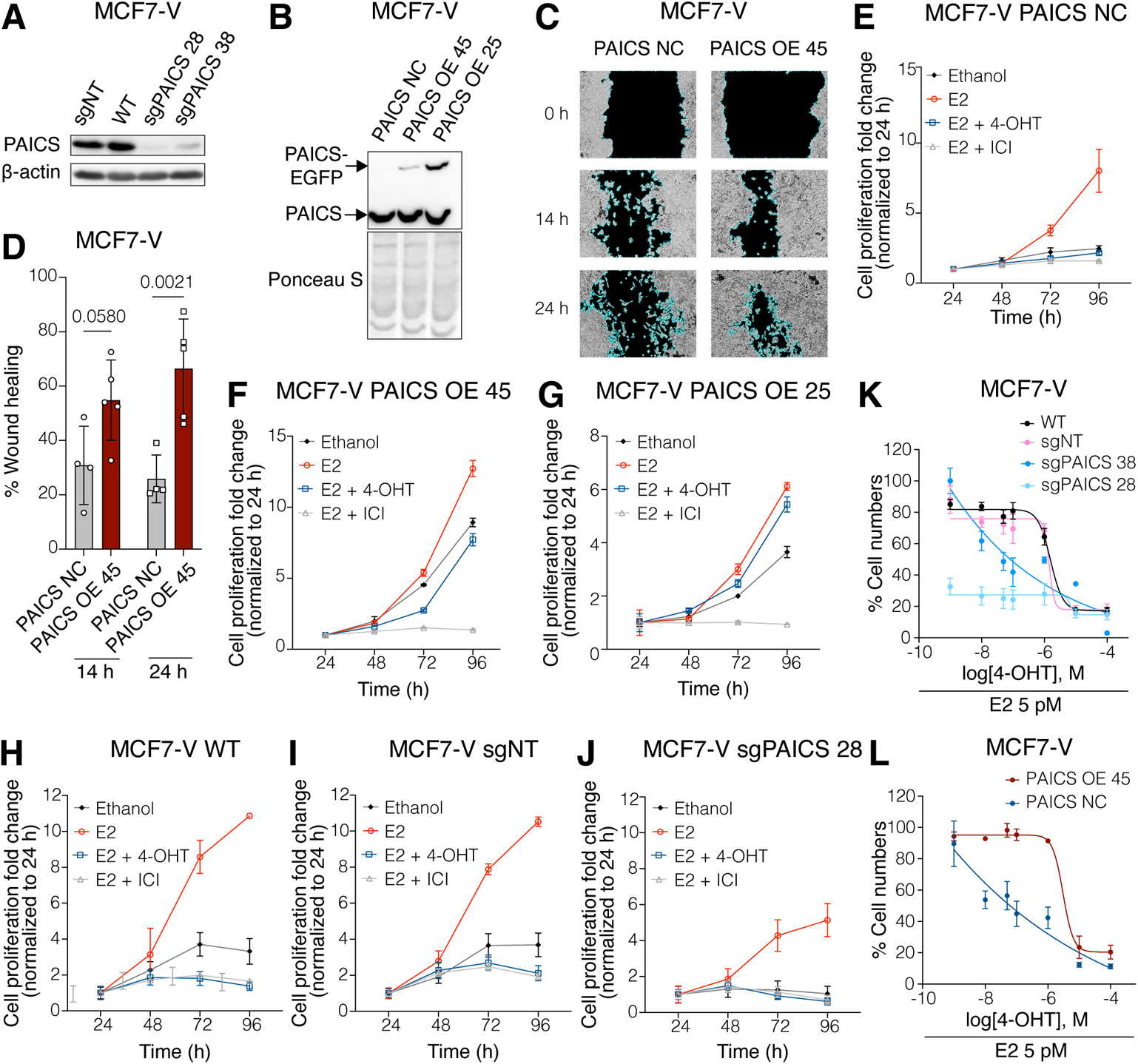
PAICS expression correlates with cell migration and estrogen-independent and tamoxifen-resistant proliferation of ERα+ breast cancer cells. **(A)** Immunoblot of PAICS showing the knockdown efficiency of the two selected *PAICS* knockdown clones sgPAICS 28 and sgPAICS 38 generated in MCF7-V cells. MCF7-V WT and sgNT (sgRPE65) cells are negative controls used interchangeably. β-actin is used as an internal control. **(B)** Immunoblot of PAICS showing overexpression by the two selected PAICS-overexpressing MCF7-V clones PAICS OE 45 and PAICS OE 25. A transduced clone that did not show PAICS overexpression (PAICS NC) is used as a negative control. The PAICS-EGFP fusion protein is detected at about 77 kDa. Ponceau S staining is used as a protein loading control. **(C)** Cell migration assay of MCF7-V cells without and with PAICS overexpression. Representative phase-contrast images were captured at the time of the scratch (0 h), after 14 h and 24 h. Images for all replicates (including these) are presented in fig. S3F. **(D)** Bar graph showing the % wound healing of MCF7-V cells without and with PAICS overexpression. Wound areas were analyzed with the software ImageJ. The data are represented as means ± SD of at least 3 independent experiments. The statistical significance between the groups was analyzed by two-way ANOVA, and *p*-values < 0.05 were considered statistically significant. **(E** to **J)** Proliferation of the indicated MCF7-V clones treated with either vehicle (ethanol), E2 (100 pM), E2 (100 pM) + 4-OHT (100 nM), or E2 (100 pM) + ICI (100 nM), measured with a crystal violet assay. The data are represented as means ± SD of at least 3 independent experiments. **(K** and **L)** Dose-response curves with increasing doses of 4-OHT with sgPAICS 28 and sgPAICS 38, WT, and sgNT cells (K), and PAICS OE 45 and PAICS NC cells (L), measured by a crystal violet assay. For each curve, a vehicle-treated control group was set to 100%. The data are represented as means ± SD of at least 3 independent experiments.

To evaluate the effect of the differential protein expression levels of PAICS on the response to 4-OHT, we treated wild-type untransfected (WT) MCF7-V cells, a pool of sgNT clones, the two clones that are deficient in PAICS, the two clones overexpressing PAICS, and PAICS NC cells with increasing doses of 4-OHT. Knockdown of PAICS resulted in an increased sensitivity to 4-OHT as compared to either WT or sgNT cells (Fig. 3K). By comparing the approximate concentration required for 50% inhibition (IC50), we found that PAICS deficiency resulted in an enhanced potency of 4-OHT by about 12- and 4,000-fold in sgPAICS 38 and sgPAICS 28 knockdown clones, respectively, as compared to sgNT cells (Fig. 3K). Conversely, overexpression of PAICS led to a 4-OHT-resistant phenotype when compared to PAICS NC cells (Fig. 3L).

Overexpression of PAICS reduced the potency of 4-OHT by about 50-fold in PAICS OE 45 cells as compared to PAICS NC cells (Fig. 3L). Hence, PAICS expression levels are directly correlated with resistance of ERα+ breast cancer cells to 4-OHT treatment.

### Interplay between PAICS and ERα

Since we had found that increased PAICS expression levels promote estrogen-independent proliferation of MCF7-V and T47D cells (Fig. 3, E to G, and fig. S3, I to K), and resistance to 4-OHT (Fig. 3, E to G, K, and L, and fig. S3, I to K), we wondered whether PAICS affects expression of ERα, and what role it plays in ERα signaling and tamoxifen resistance. We determined ERα expression at the mRNA (*ESR1*) and protein levels and found that PAICS deficiency in MCF7-V cells resulted in an increase in the basal *ESR1* mRNA and ERα protein levels as compared to parent MCF7-V cells (Fig. 4, A to C). This increase of ERα levels is associated with an increase in the basal and E2-induced transcription of the ERα target genes *CXCL12*, *TFF1*, and *XBP1* (Fig. 4, D and E, and fig. S4A). In contrast, overexpression of PAICS significantly diminished *ESR1* mRNA and ERα protein levels in MCF7-V (Fig. 4, A and B, and fig. S4A). Similarly, minimal ERα protein levels were observed in T47D when PAICS was overexpressed (fig. S4B). This downregulation was also associated with a diminished basal and E2-induced transcription of the ERα target genes *CXCL12*, *TFF1*, and *XBP1* in MCF7-V (Fig. 4, F and G, and fig. S4A). To verify that the reduction of ERα levels by PAICS overexpression is not due to a non-specific clonal effect, we transfected PAICS OE 45 and PAICS OE 25 cells with an expression vector for sgPAICS. Indeed, the depletion of both endogenous and exogenous PAICS upregulated ERα protein levels (fig. S4C). These results demonstrate that there is an inverse correlation between PAICS and ERα expression at both the mRNA and protein levels.

**Fig. 4.**
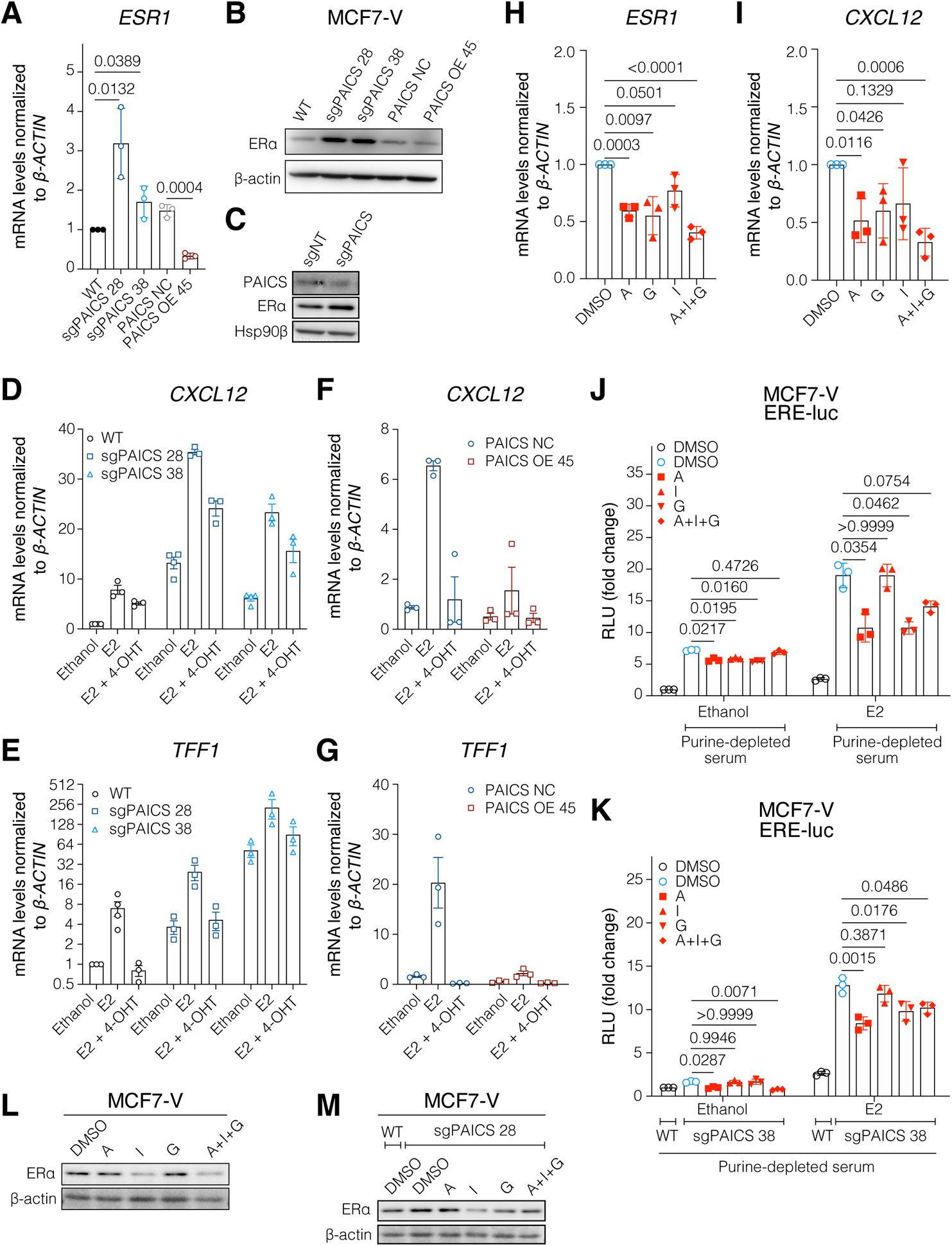
Interplay between PAICS and ERα. **(A)** mRNA levels of the ERα gene *ESR1* measured by RT-qPCR in MCF7-V cells as indicated. Ct values were first normalized to *β-ACTIN*, then values corresponding to WT were set to 1. **(B)** Immunoblot of ERα in total cell lysates. **(C)** Immunoblots of ERα and PAICS in total cell lysates of MCF7-V cells transiently transfected as indicated. Hsp90β was used as loading control. **(D** to **G)** mRNA levels of the ERα target genes *CXCL12* and *TFF1* in indicated MCF7-V cells; E2 was used at 1 nM. For panels D and E, values corresponding to WT treated with vehicle were set to 1. For panels F and G, values corresponding to PAICS NC treated with vehicle were set to 1. **(H** and **I)** mRNA levels of *ESR1* and *CXCL12* in WT MCF7-V cells starved in medium containing purine-depleted serum for 72 h and then treated with either DMSO (vehicle), A (5 µM adenosine), G (5 µM guanosine), I (5 µM inosine), or a combination of the 3 nucleosides (A + I + G), each at 5 μM. Values corresponding to vehicle were set to 1. **(J)** Luciferase reporter assay for endogenous ERα activity in MCF7-V WT cells transiently transfected with the ERE-Luc reporter plasmid. The relative luciferase activities (RLU) are relative to the activities of the internal transfection standard Renilla luciferase. Cells were starved in medium containing purine-depleted serum for 72 h and then treated as indicated. In parallel, a control group of cells that were not purine-starved were incubated in medium containing purine-rich serum for 72 h and then treated with DMSO (with or without E2). Luciferase activity of the control group incubated in purine-rich medium and treated with the vehicle (ethanol) was set to 1. **(K)** Luciferase reporter assays in sgPAICS 38 cells using the ERE-Luc reporter plasmid. Cells were treated as in panel J. In parallel, a control group of non-transduced WT MCF7-V cells were purine-starved and then treated with DMSO (with or without E2). Luciferase activity of the MCF7-V WT control group treated with ethanol was set to 1. **(L)** Immunoblot of ERα in total cell lysates of MCF7-V WT cells, hormone-starved for 72 h and then treated with either DMSO, A (100 μM), I (100 μM), G (100 μM), or a combination of the 3 nucleosides (A + I + G) each at 100 μM. **(M)** Immunoblot of ERα in sgPAICS 28 cells, treated as in panel L. For bar graphs, data are represented as means ± SD of at least 3 independent experiments. For panels A, H, I, J, and K, statistical significance between the groups was analyzed by unpaired Student’s t-tests (A, H, and I) or two-way ANOVA (J and K), and *p*-values < 0.05 were considered statistically significant.

Considering the biochemical function of PAICS, it might affect ERα expression through its impact on purine biosynthesis. To explore the mechanism, we treated WT MCF7-V cells cultured in medium with purine-depleted serum with either adenosine, guanosine, inosine, or a combination thereof. We observed that the expression of *ESR1* itself and of the ERα target genes *CXCL12*, *GREB1*, and *XBP1* is reduced by added adenosine and guanosine nucleosides, but less consistently by inosine (Fig. 4, H and I, and fig. S4, D and E). We also determined the impact of purines on ERα transcriptional activity with a luciferase reporter assay using the estrogen response element-luciferase reporter ERE-Luc. Incubation with medium containing purine-depleted serum was associated with an induction of basal and E2-induced ERα activity, and the addition of nucleosides reduced activity to varying extents (Fig. 4J). Remarkably, a very similar pattern could be seen upon depletion of PAICS (Fig. 4K). The addition of the three nucleosides, notably in combination, caused the downregulation of ERα in both WT and PAICS-deficient MCF7-V cells (Fig. 4, L and M, and fig. S4F). Hence, both ERα expression and activity are affected by the availability of purine nucleotides.

We speculated that the upregulation and increased ERα availability and activity in the case of a PAICS deficiency could sensitize cells to 4-OHT, and that PAICS overexpression could have the opposite effect. To confirm our hypothesis, we simulated the effect of a PAICS deficiency on ERα levels by overexpression of full-length ERα in MCF7-V cells (fig. S4G). A dose-response analysis of ERα-overexpressing clones with 4-OHT showed a significant enhancement of potency by about 9-fold and 16-fold for ERα overexpression clones OE 17 and OE 24, respectively (fig. S4, H and I). Therefore, PAICS expression is a determinant of ERα mRNA and protein expression and influences the cellular responses to ERα agonists and inhibitors, such as tamoxifen.

### Increased expression of PAICS correlates with cAMP/PKA activation in ERα+ breast cancer cells

Since PAICS deficiency would impair the synthesis of purine nucleotides, including adenosine monophosphate (AMP), we expected a decrease in the pool of ATP and hence, a reduction in cAMP levels. By measuring the basal ATP levels in PAICS-deficient (Fig. 5A) and - overexpressing MCF7-V cells (Fig. 5B), we found that PAICS expression levels are directly correlated with ATP levels (Fig. 5, A and B). A similar increase in ATP was associated with PAICS-overexpression in RPE1 cells (fig. S5A). Then, we directly determined the induced levels of cAMP upon treatment with the adenylate cyclase activator forskolin and the phosphodiesterase inhibitor 3-isobutyl-1-methylxanthine (IBMX), a cocktail hereafter referred to as FI. In the case of a PAICS deficiency, both the peak and the cAMP levels after 6 hours were lower than in the sgNT control cells (fig. S5B). Conversely, the overexpression of PAICS resulted in higher sustained levels of cAMP (fig. S5C). Modulation of cAMP levels would affect the extent of PKA activation and the phosphorylation of its downstream substrates, including the transcription factor CREB1. Activation of CREB1 by phosphorylation is known to contribute to increased growth and survival of cancer cells (*41, 42*). We determined PKA activity using the PKA-SPARK reporter (*43*). This system enabled us to visualize PKA activity as GFP-based phase separation droplets. A significant reduction in PKA activity at both the basal and FI-induced levels was observed in PAICS-deficient cells compared to WT (Fig. 5C and fig. S5D). To assess the phosphorylation of the key PKA substrate CREB1, FI-induced CREB1 phosphorylation was visualized by immunoblotting with a phosphoserine-specific antibody. We found that PAICS deficiency was associated with reduced CREB1 phosphorylation upon treating cells with FI (Fig. 5D and fig. S5E). Overexpression of PAICS augmented both the initial and the sustained phosphorylation of CREB1 (Fig. 5E). In a luciferase reporter assay for CREB1, deficiency of PAICS led to a significant reduction in cAMP response element (CRE) activity both at the basal and FI-induced levels (Fig. 5F). Therefore, PAICS expression determines the levels of ATP, the induced levels of cAMP, and the activity of PKA and its substrates.

**Fig. 5.**
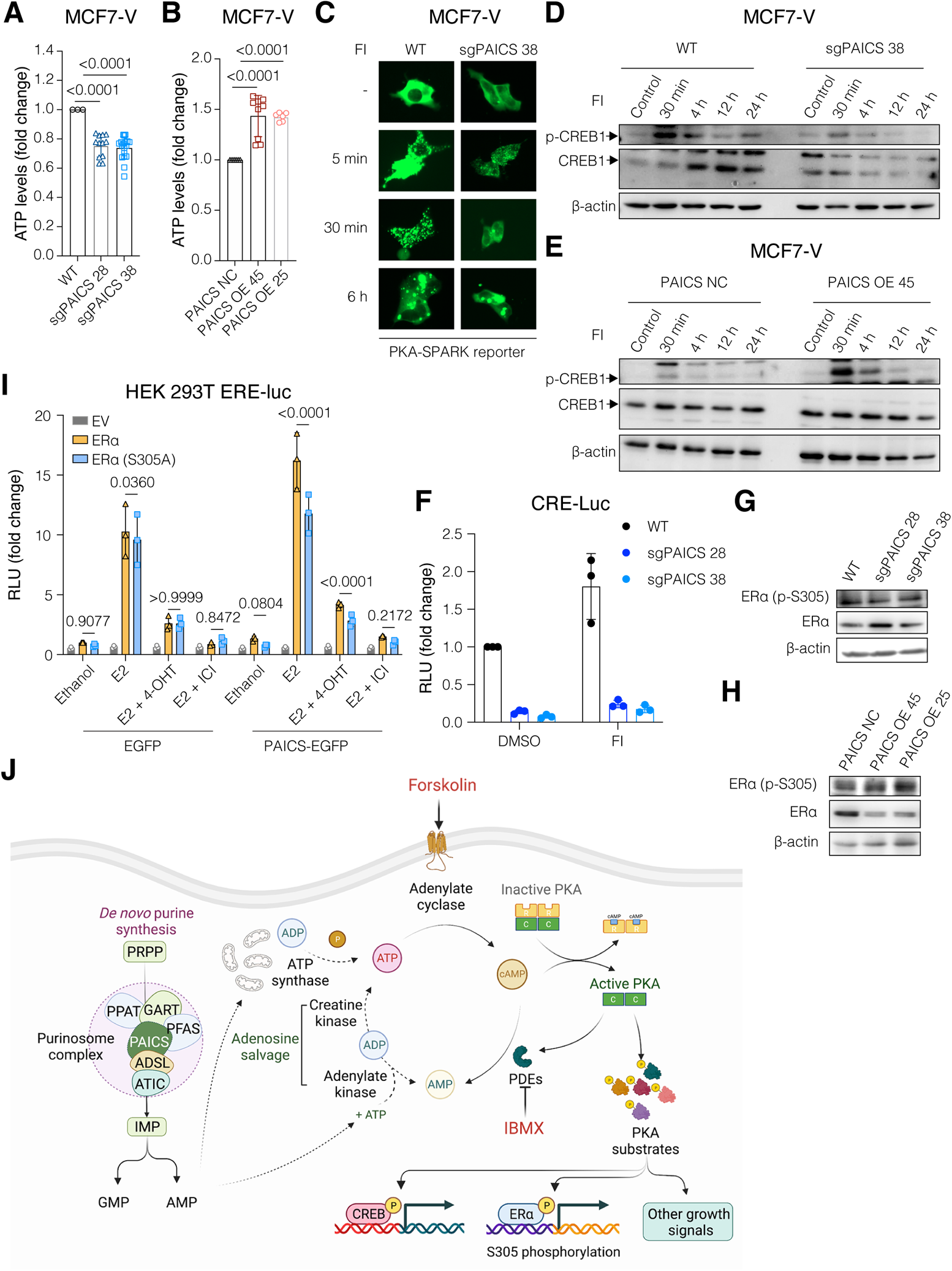
Increased expression of PAICS correlates with activation of cAMP/PKA signaling in ERα+ breast cancer cells. **(A** and **B)** ATP levels measured using CellTiter-Glo reagent of equal numbers of cells. Luminescence signal of MCF7-V WT (A) or PAICS NC (B) was set to 1. **(C)** Representative images of cells transiently transfected with the PKA-SPARK reporter plasmid. Cells were treated with FI for 5 min, 30 min, and 6 h to induce cAMP/PKA signaling. Green-fluorescent phase separation droplets are indicative of active PKA. Images for all replicates (including these) are presented in fig. S5D. **(D** and **E)** Immunoblots of phosphorylated (S133) and total CREB1 in total cell lysates. Cells were treated with FI for 30 min, 4 h, 12 h, and 24 h to induce the phosphorylation of CREB1 downstream of active PKA. **(F)** Luciferase reporter assay for endogenous CREB1 activity in the indicated MCF7-V cells transiently transfected with the CRE-Luc reporter plasmid. Cells were hormone-starved for 72 h and then treated with either vehicle (DMSO) or FI. Luciferase activities of MCF7-V WT cells treated with vehicle were set to 1. **(G** and **H)** Immunoblots of phosphorylated (S305) and total ERα. **(I)** Luciferase reporter assay for exogenously expressed ERα in HEK 293T cells transiently co-transfected with the ERE-Luc reporter, and either the plasmid PAICS-EGFP for PAICS overexpression or the empty vector pEGFP-C1 (EGFP) as a control. In addition, transfection mixtures contained either empty vector pSG5 (EV), HEG0 (ERα), or the corresponding plasmid for expression of ERα S305A. The relative luciferase activities (RLU) are relative to the activities of the internal transfection standard Renilla luciferase. Cells were hormone-starved for 72 h and then treated with either ethanol, E2 (1 nM), or E2 (1 nM) + 4-OHT (100 nM), or E2 (1 nM) + ICI (100 nM) for 24 h. Luciferase activities of the control cells transfected with ERα and treated with vehicle was set to 1. **(J)** Schematic illustration summarizing the influence of PAICS levels on cAMP/PKA signaling. PDEs, phosphodiesterase enzymes. For bar graphs, data are represented as means ± SD of at least 3 independent experiments. For panels A, B, and I, statistical significance between the groups was analyzed by unpaired Student’s t-tests (A and B) or two-way ANOVA (I), and *p*-values < 0.05 were considered statistically significant.

Activation of cAMP/PKA promotes ERα activity both in the presence and in the absence of estrogen or tamoxifen. Phosphorylation of ERα by PKA at different sites, including S305, contributes to its activation and tamoxifen resistance (*15, 16, 44*). Therefore, we speculated that overexpression of PAICS might promote the phosphorylation of S305 of ERα, which might contribute to the observed estrogen-independent growth and resistance to tamoxifen (Fig. 3, E to G, K, and L, and fig. S3, I to K). Immunoblotting for ERα phospho-S305 showed that PAICS deficiency is associated with reduced phosphorylation of ERα at S305, whereas the overexpression of PAICS correlates with hyperphosphorylation at this site. This is particularly obvious when one considers the fact that total ERα levels are inversely affected by PAICS (Fig. 5, G and H, and fig. S4B).

Further validation using an ERE-Luc reporter assay in human embryonic kidney HEK 293T cells revealed that overexpression of PAICS shifted the activity of ERα to be more dependent on S305, since the E2-induced transcriptional activity of the S305A mutant was reduced compared to that of wild-type ERα (Fig. 5I). These results are consistent with the conclusion that the expression of PAICS is directly correlated with phosphorylation of ERα at S305, which contributes to tamoxifen resistance (*15, 16, 44*) (Fig. 5J).

### Expression of PAICS is correlated with mTOR activity and the global rate of protein translation

Nucleotide sensing stimulates mTOR signaling through the regulation of the TSC complex and the GTP-binding protein Rheb (*45*). We therefore expected that PAICS protein levels might be correlated with mTOR activity. By immunoblotting, we found that PAICS deficiency correlates with decreased phosphorylation of mTOR at S2448 and of its downstream target S6, and increased phosphorylation of the translation initiation factor eIF2α (Fig. 6A). Reduced and increased phosphorylation of S6 and eIF2α, respectively, lead to reduced translation. Conversely, the overexpression of PAICS is correlated with an increase in mTOR activity, S6 phosphorylation, and decreased phosphorylation of eIF2α (Fig. 6B). When PAICS was overexpressed in non-cancerous RPE1 cells, we also detected increased phosphorylation of mTOR and NFκB, a downstream target of the mTOR signaling pathway (fig. S6A). To confirm that the reduced activity of mTOR in PAICS-deficient cells is due to a depletion of purine nucleotides, the phosphorylation level of NFκB was used as a readout in response to the addition of purine nucleosides. We noticed a reduction in NFκB phosphorylation upon purine depletion of MCF7-V cells and this reduction could be reversed by supplementing the medium with the nucleosides adenosine, guanosine, and inosine (fig. S6B). Similarly, nucleosides reverted the reduced activation of mTOR and phosphorylation of its substrate 4E-BP1 in PAICS-deficient cells (fig. S6C). Phosphorylation of 4E-BPs, including 4E-BP1 prevents the sequestration of the initiation factor eIF4E, thereby promoting translation. Increased expression, availability, and phosphorylation of eIF4E, and hence global translation, was reported to be associated with tamoxifen resistance (*46, 47*). Differential regulation of mTOR activity and eIF2α implies a direct correlation between PAICS and the global protein translation machinery. To test this, PAICS-overexpressing cells were treated with puromycin for specific time points and tested for the incorporation of puromycin into nascent polypeptides by immunoblotting with an anti-puromycin antibody. We observed more puromycin incorporation when PAICS was overexpressed and hence, increased translation (Fig. 6C). Therefore, PAICS levels are directly correlated with mTOR activation and global translation.

**Fig. 6.**
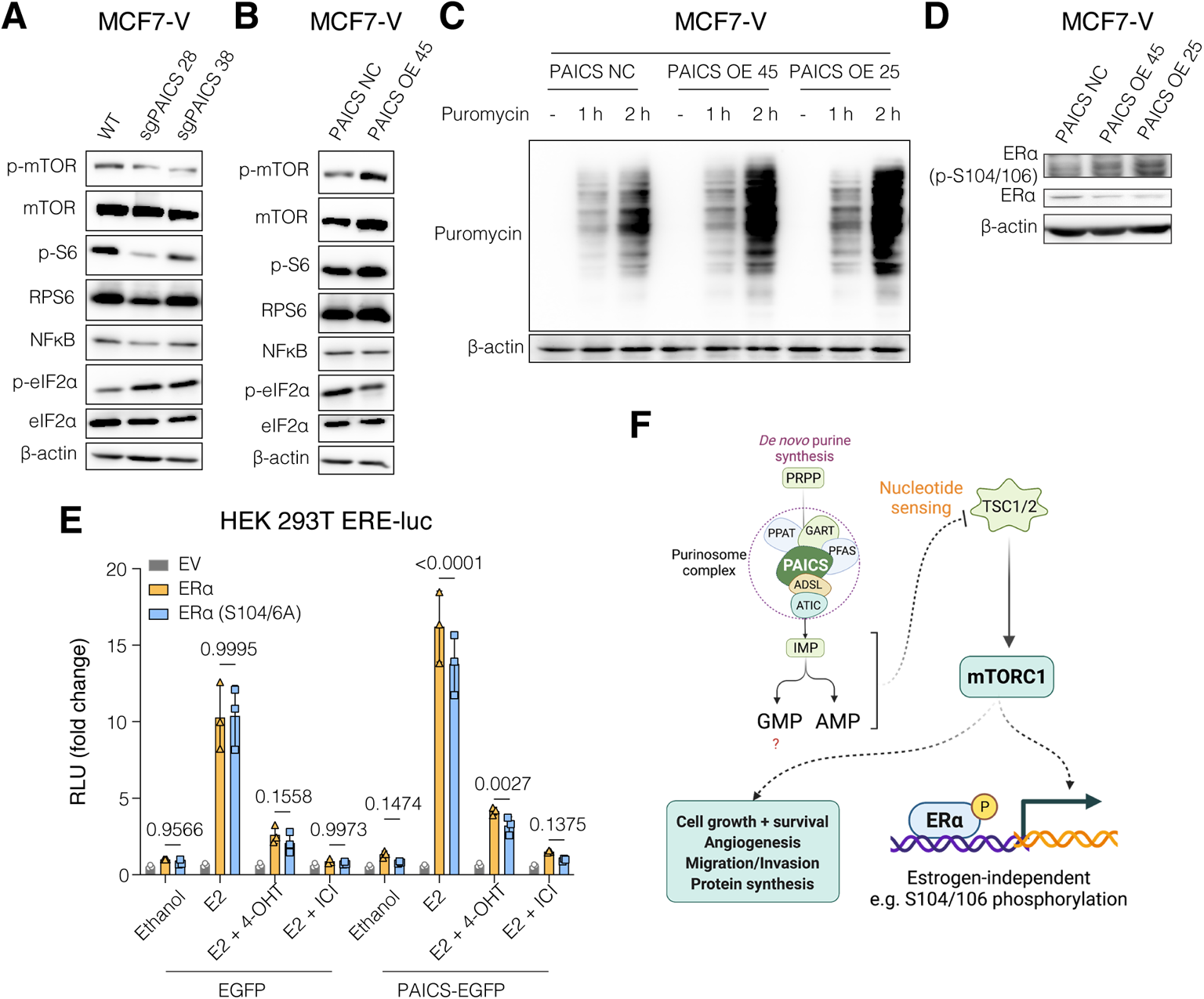
Expression of PAICS is correlated with mTOR activity and the global rate of protein translation. **(A** and **B)** Immunoblot of the indicated proteins. **(C)** Immunoblot of puromycin-labelled nascent polypeptides in total cell lysates. Cells were treated without or with puromycin (1 μM) for 1 h and 2 h. **(D)** Immunoblot of phosphorylated S104/106 of ERα and total ERα. **(E)** Luciferase assays in HEK 293T cells using the ERE-Luc reporter, as in Fig. 5I. Data are represented as means ± SD of at least 3 independent experiments. Values for EV and ERα are the same as those in Fig. 5I as both S305A and S104/6A were tested in parallel. The statistical significance between the groups was analyzed by two-way ANOVA, and *p*-values < 0.05 were considered statistically significant. **(F)** Schematic illustration summarizing the influence of PAICS levels on mTOR signaling.

Activation of Akt/PI3K/mTOR can promote the activation of ERα by estrogen and tamoxifen, and in a ligand-independent manner (*48*). In the presence of estrogen, mTOR was previously shown to boost ERα activity through the phosphorylation of S104 and S106, which are located in AF-1 (*49*). S104/106 were shown to be phosphorylated in a ligand-independent manner by MAPK *in vitro* and *in vivo* and their phosphorylation is required for the agonistic effects of tamoxifen (*50*). Therefore, we expected that PAICS expression might influence ERα activity through mTOR-stimulated phosphorylation of S104/106. We observed an increased phosphorylation at S104/106 when PAICS was overexpressed, and again most obviously when one considers the reduced levels of total ERα (Fig. 6D and fig. S4B). Additionally, using an ERE-luciferase reporter assay with HEK 293T cells, we found a significant impairment of the transcriptional activity of the ERα double mutant S104/106A upon PAICS overexpression, which implies that the activity of ERα is shifted to be more dependent on these sites (Fig. 6E). Taken together, we conclude that PAICS expression levels are directly correlated with mTOR activity, global translation, and phosphorylation of ERα at S104/106, which can all contribute to tamoxifen resistance.

### Pharmacological inhibition of PAICS sensitizes ERα+ breast cancer cells to 4-OHT

Since PAICS levels can be upregulated in cancer (Fig. 2, A to C, and fig. S2, A to D), we sought to evaluate PAICS as a potential drug target and to test the efficacy and the safety profiles of the small-molecule PAICS inhibitor MRT00252040 (MRT) (*45, 51*). We first generated dose-response curves of MRT for the ERα+ breast cancer cell lines MCF7, T47D, and MCF7-V, the tamoxifen-resistant cell lines MCF7/LCC2 and MCF7/TamR (*52, 53*), and for the non-cancerous breast epithelial cell line MCF10A (fig. S7A).

Calculations of approximate IC50 values showed that MRT is slightly more potent in the tamoxifen-resistant cell lines MCF7/LCC2 and MCF7/TamR by about 4- and 1.7-fold, respectively, by comparison with the tamoxifen-sensitive wild-type MCF7 cells (fig. S7B). Interestingly, MCF10A showed a much lower sensitivity than the cancerous cells. The IC50 values of MCF7 (about 460 nM) and MCF10A (about 110 μM) cells indicate a large therapeutic window of up to 240:1 (fig. S7B). To determine whether MRT treatment would result in an increased 4-OHT sensitivity of ERα+ breast cancer cells, two doses of MRT corresponding to ∼IC10 (50 nM) and ∼IC20 (120 nM) (fig. S7C) were combined with increasing doses of 4-OHT with our panel of cell lines (Fig. 7, A to F, and fig. S7, D and E). At these low doses, MRT increased the sensitivity of the tamoxifen-tolerant MCF7-V or -resistant cell lines MCF7/LCC2 and MCF7/TamR, and PAICS OE 45 cells to 4-OHT (Fig. 7, A to E, and fig. S7, D and E). Moreover, the drug combination did not result in a significant difference compared to 4-OHT alone in the non-cancerous cell line MCF10A (Fig. 7F). To further confirm the safety of the drug combination, two additional (non-breast) non-cancerous cell lines (RPE1 and BEAS-2B) were tested; similarly, the combinations did not show any additional toxicities, as compared to the treatment with 4-OHT alone (fig. S7, F and G).

**Fig. 7.**
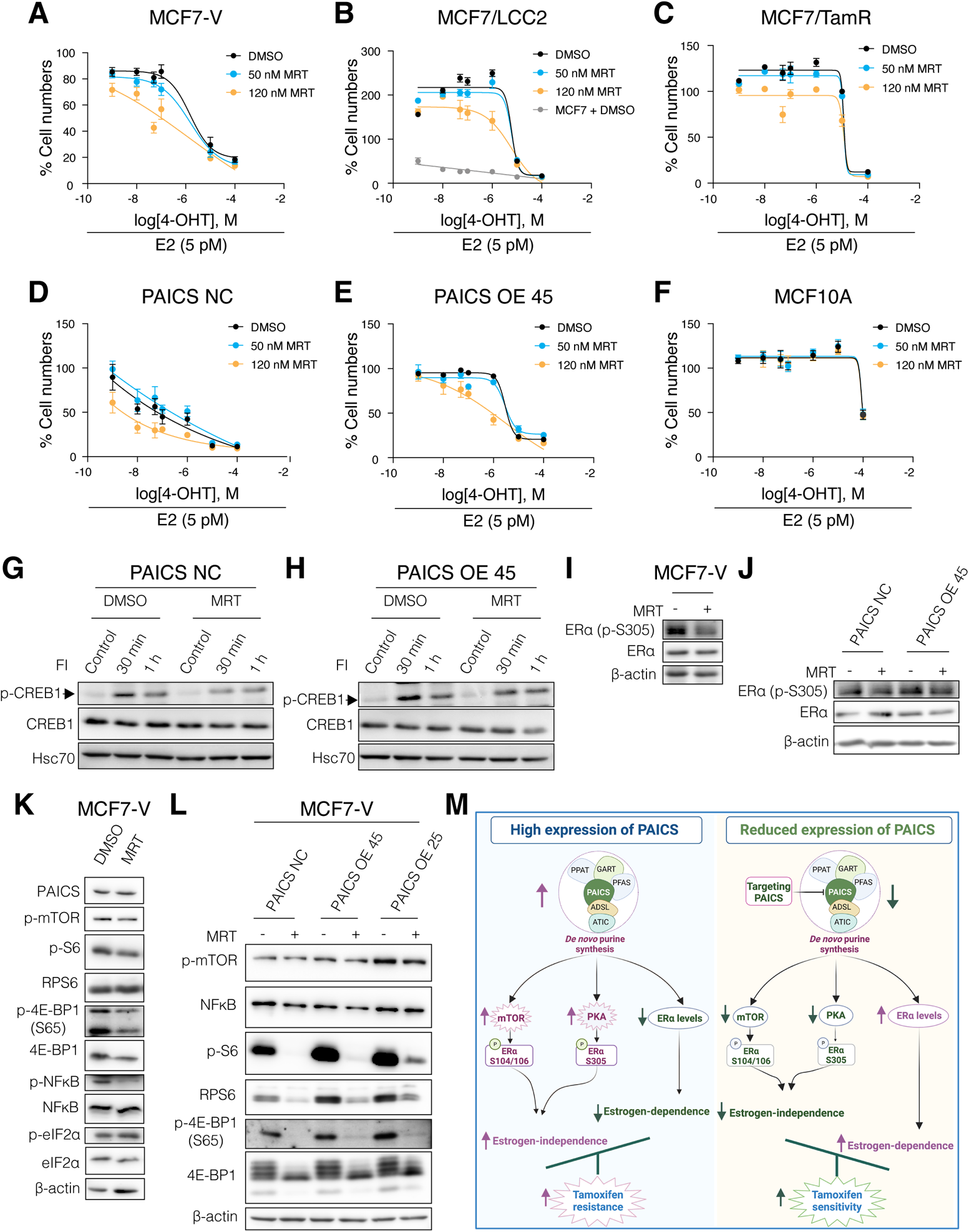
Pharmacological inhibition of PAICS sensitizes ERα+ breast cancer cells to 4-OHT. **(A** to **F)** Dose-response curves with increasing doses of 4-OHT in the presence of either vehicle (DMSO) or MRT (50 nM or 120 nM), with cell densities measured with a crystal violet assay. For each curve, a control group treated with vehicle was set to 100%. Data are represented as mean ± SD of at least 3 independent experiments. **(G** and **H)** Immunoblots of phosphorylated (S133) and total CREB1. Cells were pre-treated with either vehicle (DMSO) or MRT (500 nM) for 24 h. Then, they were further treated with FI for 30 min and 1 h. Simultaneously, untreated cells (Control) were used as a control that corresponds to basal levels of CREB1 phosphorylation. Hsc70 was used as an internal control. **(I** and **J)** Immunoblots of phosphorylated (S305) and total ERα. Cells were treated with either DMSO or MRT (500 nM) for 24 h. **(K** and **L)** Immunoblots of the indicated proteins in total cell lysates of cells treated with either vehicle (DMSO) or MRT (500 nM) for 24 h. **(M)** Schematic illustration summarizing the impact of alterations of PAICS expression levels on cell signaling and ERα activity.

We speculated based on our aforementioned results (Fig. 5 and fig. S5) that MRT might be an indirect inhibitor of cAMP/PKA signaling. Indeed, we found that pre-treatment with MRT resulted in reduced FI-induced CREB1 phosphorylation in PAICS-overexpressing MCF7-V cells (Fig. 7, G and H). MRT treatment also led to decreased phosphorylation of ERα at S305 (Fig. 7, I and J). Therefore, inhibition of PAICS with MRT causes a reduction of cAMP/PKA-induced phosphorylation of CREB1 and ERα S305.

Since we have also shown that PAICS levels correlate with mTOR activity and global protein synthesis (Fig. 6 and fig. S6), we analyzed the impact of MRT on those. We found that MRT treatment of WT MCF7-V and PAICS-overexpressing cells resulted in a reduction of the phosphorylation of mTOR, and of its substrates S6 and 4E-BP1 (Fig. 7, K and L). MRT also caused a reduction in the downstream phosphorylation of NFκB and caused the inhibitory hyperphosphorylation of eIF2α (Fig. 7, K and L). Similarly, downregulation of protein translation by MRT was observed when WT MCF7-V and PAICS-overexpressing cells were pre-treated with MRT and labelled with puromycin (fig. S7, H and I), in line with our previous findings (Fig. 6C). Therefore, the PAICS inhibitor MRT causes a reduction of mTOR activity and reduces the activity of the protein translation machinery.

## Discussion

Our work casts a new light on the role of the *de novo* biosynthesis of purines in endocrine therapy resistance of breast cancer. With a genome-wide CRISPR/Cas9 screen, we identified a set of genes that potentially contains new therapeutic targets and/or biomarkers of endocrine therapy responses (Fig. 1, B and C). One of the top hits encodes the enzyme PAICS whose role in breast cancer was largely unknown. Here we report the discovery of an important correlation of PAICS levels with tumor progression, estrogen-independent cell proliferation, and tamoxifen resistance of ERα+ breast cancer. Our findings also highlight mechanistic connections between PAICS and the cAMP/PKA and mTOR signaling pathways. From a therapeutic perspective, we propose that PAICS may be a promising drug target to overcome tamoxifen resistance of ERα+ breast cancer. Metabolic enzymes are commonly deregulated in cancer due to the increased demand caused by rapid cell proliferation (*54, 55*). The biosynthesis of purines occurs through the purine salvage or *de novo* biosynthesis pathways. The salvage pathway is predominant and involves the recycling of nucleotide metabolites to regenerate purines. The *de novo* purine biosynthesis is the biological process that converts phosphoribosyl pyrophosphate to inosine monophosphate (IMP), and from there to AMP and GMP. This process is energy-intensive and relies on ten highly conserved steps catalyzed by six different enzymes. Increased cellular demand of purines stimulates these enzymes to form a multi-component, dynamic, non-membrane bound complex known as the purinosome complex (*56*). PAICS catalyzes steps 6 and 7 of *de novo* purine biosynthesis (*57*). High PAICS expression levels are correlated with the formation of multiple tumors, including breast (*58*), bladder (*59*), prostate (*60*), gastric (*61*), colorectal (*62*), and lung (*63*) carcinomas, and acute myeloid leukemia (*51*). Hypomethylation of the *PAICS* gene was reported for lung cancer cells as compared to normal lung cells (*63*), possibly contributing to elevated expression. By comparing the expression levels of PAICS in cancerous and non-cancerous cell lines, we observed consistently higher protein levels of PAICS in the cancer setting (fig. S2, A to D). Our analysis of PAICS protein expression using a TMA of human clinical breast biopsies showed a significantly higher expression of PAICS in malignant tissue cores in both ERα+ and ERα-breast samples (Fig. 2, A to C). This could be explained by an increased dependence of cancer cells on the synthesis of the nucleoside triphosphate precursors AMP and GMP to maintain elevated rates of transcription, DNA replication, cell division, and cell signaling (*63*). Further analysis of the TMA data revealed a significant correlation of PAICS expression and breast tumor proliferation, invasiveness, grade, and TNM stage in ERα+ samples rather than ERα-ones (Fig. 2, A to G, and fig. S2, E to J). In line with this, survival analyses of ERα+ and ERα-breast cancer patients revealed a statistically significant correlation between the expression levels of *PAICS* and poor outcome in ERα+ breast cancer including those that were treated with tamoxifen. In contrast, there was no significant correlation for ERα-breast cancer (Fig. 2, H to J, and fig. S2, K to M). This partially agrees with a previous report about *PAICS* being a highly prognostic gene for breast cancer growth and aggressiveness, irrespective of ERα status (*64*). To our knowledge, a role of PAICS in ERα signaling and regulation had not been established before.

### Role of PAICS in cell migration, proliferation, and the regulation of ERα

The finding that PAICS is upregulated in tumors suggests that it might be a favorable therapeutic target. Our data show that a deficiency of PAICS results in an enhanced sensitivity of ERα+ breast cancer cells to 4-OHT, while its overexpression leads to a resistant phenotype (Fig. 3, K and L). High expression of PAICS is associated with cell proliferation, migration, and invasion (*59, 60, 62*). We show that the overexpression of PAICS in the non-cancerous cell line RPE1 is associated with increased cell migration (fig. S3, A to D). Moreover, in ERα+ breast cancer cells, PAICS was found to enhance cell migration and to modulate cell proliferation towards estrogen-independence (Fig. 3, C to J, and fig. S3, F to K). The observed increased cell migration is consistent with an *in vivo* animal study showing that PAICS depletion could inhibit primary and metastatic breast tumor growth (*64*). In the same study, PAICS depletion led to a more pronounced reduction in cell proliferation *in vivo* than *in vitro*. Differential effects of 4-OHT and ICI are notable in the cell proliferation profiles of ERα+ breast cancer cells that overexpress PAICS (Fig. 3, E to G, and fig. S3, I to K). Enhancement of cell proliferation in the absence of estrogen and in the presence of 4-OHT would imply either an upregulation in signaling pathways that promote cell growth and proliferation in general or an altered mode of activation of ERα. Estrogen-independent or 4-OHT-resistant, or even 4-OHT-promoted activation of ERα might occur through AF-1 or through the effects of phosphorylation of S305 in the hinge domain, although a contribution of AF-2 cannot be excluded (*4, 10, 15–17, 44, 49, 50, 65*). The inhibition of the PAICS-stimulated proliferation by ICI demonstrates that the cells are still dependent on ERα. In contrast, the depletion of PAICS sensitized the cells to both 4-OHT and ICI and restricted the proliferation of cells to the presence of estrogens. These findings suggest that PAICS expression affects the regulation of ERα activity and the response of ERα to different ligands.

Expression of ERα is an essential determinant of the tamoxifen response. We demonstrated that there is an inverse correlation between PAICS levels and the expression levels of ERα mRNA and protein (Fig. 4, A to G, and fig. S4, A to C). Independent of the levels of PAICS, we found that overexpression of ERα is sufficient to enhance the sensitivity of breast cancer cells to 4-OHT treatment (fig. S4, G to I). Taken together, these results suggest that PAICS levels affect the expression of *ESR1* (ERα) at the transcriptional level and that this contributes to the modulated response to 4-OHT. This is consistent with previous studies showing that low levels of ERα expression can lead to tamoxifen resistance (*46, 66, 67*). ERα expression can be regulated by several transcription factors, DNA methylation, histone modifications, RNA-binding proteins, and miRNAs (*46*). The exact mechanisms of how PAICS or purines regulate *ESR1* expression remain to be elucidated. An interplay between ERα and *de novo* purine biosynthesis is also supported by a previous report showing that ERα signaling induces a metabolic switch in breast cancer cells, including increased biosynthesis of purines (*68*). In the same study, *PAICS* and *PPAT*, which encodes another enzyme of the *de novo* purine biosynthesis pathway, were identified as ERα target genes. The downregulation of the transcription of *ESR1* at higher levels of PAICS might be due to a regulatory, perhaps even negative feedback role of purines on the transcription of *ESR1*, or due to hypothetical purinosome-independent functions of PAICS. A previous report has shown that the availability of purines and purine nucleotides can regulate the transcription of a broad class of genes that are lacking specific consensus sequences proximal to their promoters (*69*). It remains to be seen whether *ESR1* falls into this category.

### PAICS levels determine cAMP/PKA activation and ERα phosphorylation at S305

Apart from the crucial necessity of purines for the generation of DNA and RNA molecules, purine nucleotides such as ATP and GTP are indispensable for providing cellular energy and for signaling (*70*). Depending on the energy needs, macromolecules such as carbohydrates, proteins, and fatty acids can be utilized by mitochondrial and glycolytic pathways to fuel the cells with ATP. This requires a sufficient supply of its cytosolic precursor adenosine diphosphate (ADP). ADP in turn results from the phosphorylation of AMP, which is synthesized from IMP through a two-step enzymatic reaction. Our results revealed that PAICS expression levels can impact the cellular pool of ATP (Fig. 5, A and B, and fig. S5A). This might result from increased synthesis of its precursor IMP by the *de novo* purine biosynthetic pathway. This possibility ties well with a recent study showing a reduction in the levels of AMP, ADP, and ATP upon using chemical inhibitors of purine biosynthesis (*45*). Another speculation would be that PAICS or the purinosome might have a role in the regulation of mitochondrial function. An interplay between mitochondria and the purinosome is very plausible since recent studies using super-resolution microscopy revealed their colocalization (*71, 72*). Moreover, the multistep process of the *de novo* biosynthesis of purines requires a coordinated supply of energy, substrates, amino acids, and co-factors from mitochondria (*71, 72*).

The levels of ATP and the capacity of its synthesis might impact directly or indirectly multiple cellular signaling pathways (*73*). Upon activation of the enzyme adenylate cyclase, ATP is converted to the second messenger cAMP. cAMP-activated PKA can drive tumor formation in multiple endocrine tissues, including breast (*74–76*). An inactivating mutation of the *PRKAR1A* gene, which encodes the regulatory subunit R1α of PKA, was shown to increase PKA activity and to promote carcinogenesis in mammary tissues (*74*). Downregulation of RIα and PKA overexpression were shown to be associated with increased phosphorylation of S305 of ERα and tamoxifen resistance in breast cancer (*16*). Clinical reports showed that phosphorylation of S305 of ERα was associated with resistance to tamoxifen adjuvant therapy in premenopausal women (*16, 77, 78*). Previous work from our laboratory had shown that, in response to cAMP, the phosphorylation of the coactivator-associated arginine methyltransferase 1 (CARM1) at S448 by PKA is indispensable for the activation of unliganded ERα, albeit insufficient on its own (*79*). Later on, other ERα interactors were discovered to contribute to the activation of ERα in response to cAMP, notably LSD1, Co-REST, HDAC1/6, Hsp90 (*80*), and more recently CREB1 (*81*). Here we showed that PAICS levels were directly correlated with FI-induced cAMP levels, PKA activity, and the phosphorylation of PKA substrates, such as S133 of CREB1 and S305 of ERα (Fig. 5, C to J, and figs. S4B, and S5, B to E). Regulation of ERα activity by PKA is specific to tissue, cell type, and target gene. Although PKA promotes ligand-independent activation of ERα, it was also known to regulate ERα activity in the presence of estrogen (*82–84*), and notably to decrease the estrogen dependence of ERα activation for specific genes in MCF7, T47D, and MCF7/LCC2 cells (*11*). Moreover, it was reported to cause a switch of the behavior of tamoxifen from antagonist to agonist due to an arrest of ERα in an active conformation that is able to recruit coactivators. This was found to convert tamoxifen to a growth stimulator rather than an inhibitor when S305 of ERα is phosphorylated (*16, 65*). Therefore, we propose that the impact of cAMP/PKA signaling on ERα activity explains how PAICS levels can be linked to estrogen-independent and 4-OHT-resistant proliferation of ERα+ breast cancer cells.

### PAICS correlates with mTOR activity and ERα phosphorylation at S104/106

We also explored the role of the PI3K/Akt/mTOR signaling pathway, which is known to regulate cell survival, proliferation and metabolic homeostasis (*20*). PI3K represents the most commonly altered pathway in breast cancer and a critical hub in hormone-independence and endocrine therapy resistance of ERα+ breast cancer (*48, 85*). Activation of PI3K has been shown to be associated with poor recurrence-free survival of breast cancer patients after tamoxifen adjuvant therapy (*85*). A crosstalk between PI3K/Akt/mTOR and ERα can lead to increased ERα transcriptional activity both in the absence of ligand and in the presence of estrogen or tamoxifen (*86, 87*). Moreover, PI3K/Akt/mTOR can induce the phosphorylation of c-Jun which is part of the AP-1 complex that contributes to ERα activity, breast cancer progression, invasiveness, and tamoxifen resistance (*88, 89*). Activation of PI3K/Akt/mTOR has been suggested to be required for the adaptation of ERα+ breast cancer to hormone deprivation (*85*). Targeting PI3K/Akt/mTOR with small-molecule inhibitors was shown to reverse hormone-independent cell growth and endocrine resistance (*85, 90*). Interestingly, data from our screens as well as those of others (*32*) highlight *TSC1* and *TSC2* as genes whose depletion boosts cell proliferation in the absence of estrogen and in the presence of 4-OHT or ICI (Fig. 1B). Our results show a strong correlation of PAICS levels with the activation of mTOR signaling and the global protein translation machinery (Fig. 6, A to C, and fig. S6A). This is supported by a recent report identifying purine nucleotides as an upstream regulatory stimulus for the activation of the mTOR complex mTORC1 (*45*). mTORC1 acutely senses alterations in the levels of adenine nucleotides by TSC and not through AMP-activated protein kinase or Rag GTPases. mTORC1 could also sense long-term changes in guanine nucleotide levels. Therefore, we speculate that the observed correlation of PAICS with mTOR activity could be explained by alterations of the levels of purine nucleotides (*45*). In agreement with this idea, our data show that supplementation with purine nucleosides rescues the reduced mTOR activity resulting from the knockdown of PAICS (fig. S6, B and C).

Phosphorylation of ERα within the AF-1 domain stimulates ERα ligand-independent activity and contributes to endocrine therapy resistance (*50*). Akt and ribosomal S6 kinase 1, which is a downstream target of mTOR, were found to promote ERα transcriptional activity by phosphorylating S167 in response to growth factors (*86, 87*). This phosphorylation is positively correlated with endocrine therapy response and better survival of patients with primary breast tumors (*67*). In response to estrogen, the mTORC1 component raptor binds to and translocates with ERα to the nucleus. This allows the interaction of mTOR with ERα and phosphorylation of S104/106, also located within AF-1 (*49*). ERα was found to be phosphorylated at S104/106 by MAPK both *in vitro* and *in vivo* (*50*), and by cyclin A-CDK2 independently of ligand (*91*). Phosphorylation at these sites is required for the agonistic activity of tamoxifen and may have a role in tamoxifen resistance (*50*). Our results indicate that the activity of ERα becomes more dependent on the phosphorylation of S104/106 when PAICS is overexpressed (Fig. 6, D and E, and fig. S4B). This shift might explain the estrogen-independent and tamoxifen-resistant cell proliferation observed in these cell lines. The phosphorylation of S104/106, when PAICS is overexpressed, might be mediated by a hyperactive mTOR, MAPK, or yet another kinase. Which kinase(s) is most relevant under these circumstances remains to be investigated.

### Pharmacological significance

Mechanistically, the small molecule PAICS inhibitor MRT has been shown to inhibit the SAICAR synthetase activity of PAICS since its binding to PAICS resulted in accumulation of metabolites upstream of SAICAR (*51*). It has been reported for leukemia that targeting PAICS using MRT reduces cell proliferation by inhibiting DNA synthesis and by promoting apoptosis *in vitro* and *in vivo* (*51*). Moreover, PAICS knockdown causes a reduction in tumor growth (*60–62*) and metastasis (*62*) in mouse xenograft models. Apart from the metabolic role of PAICS, its knockdown was found to induce DNA damage and to impair DNA damage repair by reducing HDAC1/2 deacetylase activity, and to enhance the sensitivity to DNA damaging agents like cisplatin in gastric carcinoma *in vitro* and *in vivo* (*61*). In breast cancer, targeting PAICS was shown to reduce the proliferation of breast cancer cells by inducing a cell cycle arrest (*58*). We have evaluated the therapeutic potential of MRT in ERα+ breast cancer cell lines. We demonstrate that MRT selectively affects breast cancer cell lines compared to non-malignant cells (fig. S7, A to C). We found that low doses of MRT re-sensitize 4-OHT-tolerant and -resistant cell lines to 4-OHT with minimal toxicities on non-cancerous cells (Fig. 7, A to F, and fig. S7, D to G). We therefore propose a low-dose combination of MRT + 4-OHT as a potentially non-toxic drug combination to treat tamoxifen-resistant breast cancer. We have demonstrated that MRT treatment has the potential to impair signaling by cAMP/PKA and mTOR, to reduce global protein translation and phosphorylation of S305 of ERα (Fig. 7, G to L, and fig. S7, H and I). These results provide a mechanistic explanation for how MRT could re-sensitize cells to 4-OHT and contribute to overcoming tamoxifen resistance.

Collectively, our study shows that PAICS expression levels are directly correlated with tamoxifen resistance, which can be attributed to increased cAMP-induced PKA and purine-induced mTOR signaling. cAMP/PKA and mTOR signaling might result in the phosphorylation of ERα at S305 and S104/106, respectively. Modulation of the levels of ERα phosphorylation at these sites might tune the balance of ERα activity in response to estrogen and 4-OHT. Finally, our results showed that targeting PAICS either with gene editing or with the specific small-molecule inhibitor MRT can sensitize breast cancer cells to 4-OHT (Fig. 7M).

#### Limitations

The huge number of cells required for a CRISPR/Cas9 screen made it difficult to do the screen in parallel with additional cell lines. This would have helped to confirm and to diversify the list of genetic biomarkers and/or targets. Moreover, although the inclusion of a treatment with a cell-permeable form of cAMP in the screen seemed worthwhile, we could not identify significant hits specifically related to this particular treatment. Perhaps other ways of activating PKA continuously throughout the duration of the screen might be more appropriate. Although our findings strongly suggest a correlation between PAICS and ERα expression, more studies are needed to identify the underlying molecular mechanism. Furthermore, the connection between PAICS and endocrine resistance needs to be further strengthened in the future with cohort studies of breast cancer patients who developed resistance to endocrine therapy. Finally, the small molecule PAICS inhibitor MRT is a recent investigational drug and therefore, future preclinical studies are necessary for its progression towards clinical trials.

## Materials and Methods

### Cell lines

In the following, for cell lines available from ATCC, references are given in parenthesis: MCF7 (HTB-22), BT474 (HTB-20), SKBR3 (HTB-30), MDA-MB-231 (HTB-26), HCC1937 (CRL-2336), MCF10A (CRL-10317), HEK 293T cells (CRL-3216), human skin fibroblasts CCD-1079Sk (CRL-2097), RPE1 cells (CRL-4000), BEAS-2B (CRL-9609) and BZR (CRL-9483). T47D cells were purchased from Sigma-Aldrich (#85102201). MCF7/LCC2 (*52*), and MCF7/TamR (*53*) (a gift from Denise Barrow, Cardiff University) were used as tamoxifen-resistant cell lines. RPE1 Ras, and the human skin fibroblast derivatives hTERT, hTERT LT, and hTERT LT Ras, generated as described before (*38*) were a gift from Vanessa Xavier and Jean-Claude Martinou (University of Geneva, Switzerland) (*39*). Human breast carcinoma H3396 cells (*92*) were a gift from Hinrich Gronemeyer’s laboratory (IGBMC, Strasbourg).

### Plasmids

The human Brunello CRISPR knockout pooled library (*36*) was a gift from David Root and John Doench (Addgene #73179). The vector lentiCRISPRv2 (*93*) (Addgene #52961) for expression of Cas9 was a gift from Feng Zhang (Addgene #52961). The VSV-G envelope expressing plasmid pMD2.G (Addgene #12259), and the lentiviral packaging plasmid psPAX2 (Addgene #12260) were gifts from Didier Trono’s laboratory (EPFL, Lausanne), and were used to produce lentiviral particles for expression of Cas9 and sgRNAs. For the specific knockouts of *PAICS* and *RPE65*, the selected sgRNA sequences target the 3^rd^ exon of *PAICS* and the 5^th^ exon of *RPE65* (data S5). Oligonucleotides were purchased from Microsynth (data S5), annealed, and cloned into the BbsI site of the vector lentiCRISPRv2 for expressing sgPAICS and sgRPE65. For transient transfection of sgPAICS and sgRPE65, annealed oligos were cloned into the Cas9-expressing vector pSpCas9(BB)-2A-Puro (PX459) V2.0 (Addgene #62988), which was a gift from Feng Zhang. The plasmid pPAICS-EGFP was a gift from Stephen Benkovic (Addgene #99108) and was used for the transient and stable overexpression of a PAICS-EGFP fusion protein. As a negative control for the transient overexpression of PAICS, EGFP expression plasmid pEGFP-C1 (Clontech #6084-1) was used. For the generation of ERα-overexpressing clones, the plasmid F-ER (*94*) was used. For transient expression of wild-type and mutant ERα in HEK 293T cells, plasmid HEG0 (*95*) and its derivatives HEG0 S305A and HEG0 S104/106A were used, with their expression vector pSG5 (*96*) as the empty vector control. For luciferase assays, the following reporter plasmids were used: XETL (here, referred to as ERE-Luc) (*97*), CRE-Luc (Stratagene), and the Renilla luciferase transfection control reporter pRL-CMV (Promega #E2261). To visualize PKA activity, plasmid pcDNA3-PKA-SPARK was used. It was a gift from Xiaokun Shu (Addgene #106920).

### Cell culture

MCF7/LCC2 and MCF7/TamR cells were cultured in Dulbecco’s Modified Eagle’s Medium (DMEM) without phenol red (Thermo Fisher Scientific #11880036), supplemented with 5% charcoal-treated fetal bovine serum (CHFBS), 100 μg/ml penicillin/streptomycin (Thermo Fisher Scientific #15070063), 4 mM L-glutamine (Thermo Fisher Scientific #25030081), and 100 nM 4-OHT (Sigma-Aldrich #H7904). MCF10A were cultured in DMEM/F12 medium (Thermo Fisher Scientific # 31331028) containing 5% horse serum (Bio-concept #2-05F00-I), 100 μg/ml penicillin/streptomycin, 10 ng/ml human epidermal growth factor (Sigma-Aldrich #E9644), 5 μg/ml insulin (Sigma-Aldrich #I9278), and 1 μM dexamethasone (Sigma-Aldrich #D8893). All other cell lines were cultured in DMEM/GlutaMAX medium (Thermo Fisher Scientific #31966047), supplemented with 10% FBS (PAN-Biotech #P40-37500), and 100 μg/ml penicillin/streptomycin. Before treatments, hormone deprivation was done by maintaining the cells in DMEM medium without phenol red, containing 5% CHFBS, penicillin/streptomycin, and L-glutamine for 72 h. All cell lines were maintained at 37°C with 5% CO_2_ in a humidified incubator.

### Genome-wide CRISPR/Cas9 screen

#### A. Library amplification and validation of its complexity

The protocol was adapted from the Broad Institute protocol of amplification of library plasmid DNA available from the GPP web portal (https://portals.broadinstitute.org/gpp/public/resources/protocols). Briefly, in a total of 4 cuvettes, 400 ng of plasmid library DNA were transformed into 100 μl Endura™ electrocompetent cells (Lucigen #60242-2) by electroporation with a MicroPulser (Bio-Rad, #1652100) using the Ec1 setting (1.8 kV). Within 10 seconds of the pulse, 975 μl recovery medium was added to each cuvette containing 25 μl of the transformation mixture and then transferred to a loosely capped tube containing an additional 1 ml of recovery medium. Tubes were incubated with shaking at 250 rpm for 1 hour at 37°C. The 4 tubes were pooled and mixed well. Bacteria were then spread on 5 ampicillin agar dishes of 500 cm^2^, in addition to one control dish without ampicillin, and incubated overnight at 37°C. Then, bacterial colonies were scraped into cold L-Broth medium, centrifuged, and pellets were weighed (total weight about 1 – 2 g). Plasmid DNA was purified by midi-prep using a NucleoBond Xtra Midi kit (Machery Nagel #740410.50) according to the manufacturer’s protocol. To check library representation, purified DNA was amplified by PCR using GoTaq DNA polymerase (Promega #M3001) as follows: Four parallel PCR reactions were prepared, each containing 200 ng DNA and the universal primers P5 and P7 (Table S1) used for Illumina sequencing (Microsynth) (*36*). The PCR program was: 95°C for 1 minute, 95°C for 30 seconds for DNA denaturation, primer annealing at 53°C for 30 seconds, and extension at 72°C for 30 seconds, for 28 cycles, then 72°C for 10 minutes for the final extension step. PCR amplicons were further purified using GenElute PCR Clean-Up Kit (Sigma-Aldrich #NA1020) and checked for correct sizes by agarose gel electrophoresis. Quantity was determined using a NanoDrop (Thermo Fisher Scientific). The four PCR reactions were pooled into one sample and sequenced by NGS at the iGE3 Genomics Platform of the University of Geneva using a HiSeq 4000 instrument (Illumina). Approximately 24 million single-end reads of 50 bp that contain the sgRNA sequences were obtained and checked for quality using the fastqc tool available within the Galaxy platform (https://usegalaxy.org/). Sequencing adaptors were trimmed using the cutadapt tool and the sgRNA sequences were aligned to the Brunello library index using the Bowtie tool. The amplified library contained about 90% perfectly matching sgRNAs, 0.01% of the sgRNAs were undetected, and the distribution skew ratio was 0.84 (*98*).

#### B. Preparation of lentiviral particles

The protocol was adapted from a previously described one (*98*). HEK 293T cells were seeded into eight 15 cm dishes at a density of 30 million cells/dish. The next day, each dish received a co-transfection mixture of 3.77 μg pMD2.G, 7.55 μg psPAX2, and 15.11 μg pooled library plasmid DNA in serum-free DMEM medium using PEI MAX 40K (Polysciences Inc. # 24765-100) at a DNA:PEI MAX ratio of 1:8.2. Media were changed after 12 – 14 h, and lentivirus-containing supernatants were collected after 48, 72, and 96 h after transfection. Supernatants were spun at 1000 xg for 5 min, then filtered through 0.45 μm sterile filters, and complemented with sterile 40% polyethylene glycol 8000 in 400 mM NaCl (PEG) (Sigma-Aldrich #25322-68-3) at a PEG:supernatant volume ratio of 1:3. Tubes were incubated at 4°C on a rotating wheel for 1 – 2 h and then centrifuged at 4000 xg, at 4°C for 30 minutes. The pellets containing the lentiviral particles were resuspended in fresh culture medium in one twentieth of the initial collection volume. The first two concentrated lentiviral suspensions were frozen at −80°C and only thawed on ice on the day of the third collection to be pooled into one lentiviral preparation.

#### C. Viral transduction and puromycin selection

To achieve > 500x coverage of the library plasmids for each treatment group, a total of one hundred and ten 15 cm dishes were seeded with MCF7-V cells at a density of 10 million cells/dish. 120 μl of the pooled lentiviral suspension were added per dish with 8 μg/ml polybrene (Sigma-Aldrich, #107689-10G) to enhance the transduction efficiency. The volume of concentrated viral suspension was optimized for 50% transduction efficiency in MCF7-V cells. 24 h after lentiviral transduction, cells were treated with 3 μg/ml puromycin (Cayman Chemical #13884) for 7 days. A control dish of non-transduced MCF7-V cells was used as a positive control for the puromycin selection efficiency.

#### D. Drug treatments

Following the selection of lentivirally transduced cells with puromycin, cells were trypsinized, cell pellets were resuspended in fresh culture media, and then pooled into one suspension of cells. A total of 100 million cells were collected and aliquoted into 20 tubes of 5 million cells each, centrifuged; cell pellets were washed with phosphate-buffered saline (PBS) and then kept at −20°C. These cells represented the transduced control cells before drug treatment (Control “T0”). The rest of the cell suspension was allocated into drug treatment groups, each with a total of 100 million cells. Hence, each group was represented by ten 15 cm dishes seeded with 10 million cells/dish. After 24 h, drug treatments were started, and medium with drugs was changed twice per week for a total period of 4 weeks. Treatment groups were as follows: vehicle control, E2 (100 pM), db-cAMP (10 μM), E2 (100 pM) + 4-OHT (100 nM), E2 (100 pM) + ICI (100 nM), E2 (100 pM) + 4-OHT (100 nM) + db-cAMP (10 μM), and E2 (100 pM) + ICI (100 nM) + db-cAMP (10 μM). Cells were passaged (1 into 2 plates) if any of the groups reached about 90% confluency. After treating the cells for 4 weeks, cells were harvested, and for each group, cell pellets were resuspended and pooled into one suspension of cells. Cell suspensions were aliquoted, and pellets were harvested as done for Control “T0”.

#### E. Genomic DNA extraction, PCR, and NGS

The following was adapted from published protocols (*99*). Cells were lysed in digestion buffer (100 mM NaCl, 10 mM Tris HCl pH 8, 25 mM EDTA, 0.5% SDS, and 0.1 mg/ml proteinase K), and incubated with shaking at 50°C for 14 h. Then, 1 μg/ml RNase was added to the lysates and incubated at 37°C for 1h. One volume of phenol/chloroform/isoamyl alcohol was added to the lysate, mixed well, and samples were centrifuged at 10,000 xg for 10 min. The aqueous phases at the top were transferred to new tubes, 5 M NaCl was added at 1/8 volume, and then 100% ethanol was added (twice as much as the volume of the original aqueous phase). Genomic DNA was recovered by centrifugation at 12,000 xg for 20 minutes. Pellets were rinsed twice with 70% ethanol, centrifuged at 12,000 xg for 5 minutes, and then left to air-dry for 15 minutes at room temperature. Pellets were dissolved in TE buffer at 65°C overnight or until dissolved. Genomic DNA samples of each treatment group were pooled into one sample, mechanically homogenized and sheared by passing through a syringe with a 25G needle. The size distribution of the sheared DNA was assessed by agarose gel electrophoresis, and DNA concentration quantitated using a Nanodrop. For each group, four parallel PCR reactions were prepared, each containing 4 μg of genomic DNA, the universal Illumina primer P5, and a barcoded primer P7, which was unique for each group (Table S1). PCR program set-up and purification of the PCR products were the same as for the initial validation of the complexity of the library. PCR amplicons were checked for correct sizes by agarose gel electrophoresis and DNA was quantitated using a Nanodrop. The four PCR reactions were pooled into one sample and sequenced by NGS with single-end reads of 50 bp with indices. Sequence quality was checked using the fastqc tool, sequencing adaptors were trimmed using the cutadapt tool and the sgRNA sequences were aligned to the Brunello library index using the Bowtie tool.

#### F. Data analysis

The CRISPR/Cas9 screen was done with two biologically independent replicates. Read sequences of sgRNAs were mapped to their corresponding genes (data S1) and a “β-score” was calculated using the MAGeCK-VISPR platform (*32, 100*) (data S2). “β-score” is similar to “log fold-change” and it indicates the respective enrichment (β > 0) or depletion (β < 0) of sgRNAs of a specific gene when compared to the control “T0”. For all genes, β ≥ 0.3 or β ≤ −0.3 and FDR value ≤ 0.1 or ≥ −0.1, were considered significantly enriched or depleted, respectively. We considered about 600 genes as essential for survival because their sgRNA counts were depleted in all conditions and eliminated from the subsequent analysis (data S3). Comparisons between every two groups were performed using the same cut-off values to identify significant differences between the β-scores. The counts of the 4 different sgRNAs/gene were checked in the two replicates to eliminate β-scores that were biased by sgRNA outlier values. Finally, 58 genes were identified as the top enriched or depleted genes, and β-scores were hierarchically clustered using R packages (Fig. 1B, and data S4). Literature curation narrowed down the list to 40 genes (data S5) with unknown functions in ERα+ breast cancer or endocrine therapy responses.

#### Secondary CRISPR/Cas9 knockout screen

For the validation of the effects of the knockout of the 40 genes identified with the primary screen, the knockout of each gene was done individually and subjected to the same treatment groups. One sgRNA sequence was selected from the Brunello library based on the calculated off-target score (*36*) and the consistency of read counts between the two biological replicates. sgRNA sequences corresponding to scrambled and *RPE65* gene sequences were used as controls. sgRNA oligonucleotides (data S5) were cloned into lentiCRISPRv2 as described above and previously (*93*). Preparation of lentiviruses and transduction of MCF7-V cells were done as mentioned previously for the primary screen. For the drug treatments, each knockout cell line was seeded at a density of 7000 cells/well of a 96-well plate and treated for 2 weeks with the same treatment groups as during the primary screen. Cell growth was assessed using crystal violet staining. Briefly, cells were washed with PBS, incubated with 4% formaldehyde in PBS for 20 minutes, washed again with PBS, then incubated with 0.1% crystal violet solution in distilled water for 30 minutes. The wells were washed thoroughly with distilled water and air-dried overnight. Crystals were dissolved in glacial acetic acid and the absorbance was measured at 595 nm with a Cytation 3 Image Reader (Agilent). The average % cell growth of the 40 gene knockout cell lines and the treatment groups were visualized as a heatmap using GraphPad Prism version 8.0.0 (Fig. 1C).

#### Generation of sgRNA knockout clones of PAICS and RPE65

sgPAICS and sgRPE65 lentiviral particles were produced in HEK 293T cells as described above. MCF7-V cells seeded at 60% confluency in 10 cm dishes were incubated for 48 h with complete culture medium containing 200 μl concentrated lentiviral suspensions/dish. In parallel, one 10 cm dish of MCF7-V was incubated without viral particles and used as a non-transduced control for the puromycin selection. 3 μg/ml puromycin was added to both transduced and non-transduced MCF7-V cells for 4 days or until all the non-transduced cells were dead. For the isolation of single PAICS knockout clones, cells that were resistant to puromycin were sorted into single cells and seeded into a 96-well plate at a density of 1 cell/well using a BD FACS Aria III Cell Sorter (BD Biosciences). After clonal expansion, single clones were screened for either a complete knockout or a significant knockdown of PAICS protein levels by immunoblotting with a PAICS antibody. Two clones, #28 and #38, which showed a significant decrease in the PAICS protein levels compared to WT MCF7-V cells were selected for further investigation. To generate the negative control cell line (sgNT), sgRPE65-mediated knockout cells that were resistant to puromycin were pooled.

#### Generation of stable PAICS and ERα overexpression clones

Cells were transfected with the expression vectors pPAICS-EGFP (MCF7-V, T47D, and RPE1) or F-ER (MCF7-V) using jetOPTIMUS (Polyplus #PPT117-07), according to the manufacturer’s protocol. After 48 h, cells were maintained in 1000 μg/ml G418 (Sigma-Aldrich #A1720) for 2-3 weeks or until the non-transfected control cells died completely. Individual cellular colonies were picked using 200 μl pipette tips, expanded, and analyzed by immunoblotting for the PAICS-EGFP fusion protein or ERα. For MCF7-V cells, two PAICS-overexpressing clones, #25 and #45, were selected for further validation, and clone #43, which did not show expression of exogenous PAICS, was selected as a negative control (PAICS NC). For T47D cells, clones #2 and #8 were overexpressing PAICS-EGFP, and clone #3 was selected as a negative control. For RPE1 cells, the PAICS-overexpressing clones #7, #9, and #17 were selected for further assays. Cellular morphology was visualized using an inverted light microscope. We found ERα to be overexpressed in MCF7-V clones #17 and #24, but not in clone #4, which was used as negative control (ERα NC).

#### Fluorescence immunohistochemistry of breast TMA

The human breast TMA (US Biomax, Inc. #BRF1503f) consists of 150 cores of 75 cases. It contains 3 cases of adjacent normal breast tissue, 3 breast fibroadenoma, 2 breast cystosarcoma phyllodes, 7 breast intraductal carcinoma, and 60 breast invasive ductal carcinoma cases. All cases are represented in duplicate cores. Data including TNM and pathology grade, with immunohistochemistry results for the markers HER-2/ERα/PR/Ki-67 were provided by the manufacturer (data S6). The protocol was adapted from a previously published one (*34*). First, paraffinized tissue sections were warmed up for 30 minutes at 60°C in a dry oven. Then, sections were deparaffinized and rehydrated by sequential incubation in the following solutions: xylene (2 washes, 5 minutes each), 100% ethanol (2 washes, 5 minutes each), 95% ethanol (1 wash for 5 minutes), 70% ethanol (1 wash for 5 minutes), and double distilled H_2_O (1 wash for 5 minutes). Unmasking was done by heat-induced epitope retrieval using 10 mM sodium citrate buffer pH 6. Tissues were boiled in the citrate buffer in the microwave at 600 W for 3 minutes, then left to cool until the temperature of the buffer reached 50°C, then boiled again for another 3 minutes, and left to cool to room temperature. Tissues were permeabilized and blocked in 0.3% Triton-X100 and 1% BSA in PBS (1% BSA in PBST) for 20 minutes. PAICS primary antibody was diluted at 1:25 in 1% BSA in PBST buffer, added to the tissue sections, and incubated in a humidified chamber overnight at 4°C. After two washes with PBS, tissues were incubated for 1 hour at room temperature with the anti-rabbit secondary Cy3 red-fluorescent antibody diluted to 1:1000 in 1% BSA in PBST buffer. Sections were washed twice in PBS for 5 minutes each and then nuclei were counterstained by incubation with diamidino-2-phenylindole dye (DAPI) (1:30,000 in PBS, from 1 mg/ml stock solution, Thermo Fisher Scientific #62248) for 5 minutes. Sections were washed 3 times in PBS for 5 minutes each, then rinsed for 5 minutes in a solution of 10 mM CuSO_4_, 50 mM NH_4_Cl to reduce autofluorescence, and then rinsed briefly in double-distilled H_2_O. The TMA slide was mounted using the fluorescence mounting medium Fluoromount (Sigma Aldrich #F4680) and left to dry in a dark chamber. Imaging was done using the high-throughput slide scanner Zeiss Axioscan.Z1 and Zen 2 software. Images were de-arrayed and analyzed for fluorescence intensity and area using QuPath 0.3.0 software. For each tissue section, fluorescence intensity was normalized to the fluorescent area (data S6). A threshold of fluorescence was set to 500 and values below that level were eliminated from the analysis. High resolution full scan images of the TMA slide are available upon request.

#### Cell proliferation assays

To determine the rate of cell proliferation, 25,000 cells/well were seeded into 48-well plates in DMEM without phenol red, containing 5% CHFBS, penicillin/streptomycin, and L-glutamine. After 24 h, one plate was harvested as a control before treatment (Day 0), and the rest of the plates were treated with vehicle (ethanol), E2 (100 pM), E2 (100 pM) + 4-OHT (100 nM), and E2 (100 pM) + ICI (100 nM). Plates were harvested after 2, 3, and 4 days from seeding for MCF7-V cells, and 2, 4, and 6 days for T47D cells. The proliferation of cells was quantitated by crystal violet staining (see above).

#### Wound healing assay

Cells were seeded into 6-well plates. At 100% confluency, wounds were formed by scratching the cell layers using a 200 μl pipette tip. Cells were then washed with Tris-buffered saline (TBS) and incubated in DMEM without phenol red, containing 5% CHFBS, penicillin/streptomycin, and L-glutamine. Images of specific areas of the wounds were taken at 0 h, 14 h, and 24 h after scratching. Image analysis was done by measuring the wound area using the software ImageJ. % Wound healing was calculated according to the following formula: 100 x (wound area at time 0 – wound area at time (i))/ wound area at time 0.

### Gene expression analyses

#### A. Drug treatments

Cells were hormone-deprived before treatments for 72 h and seeded at a density of 200,000 cells/well of a 6-well plate. Then, cells were treated with ethanol, E2 (1 nM), and E2 (1 nM) + 4-OHT (100 nM) for 24 h. Assays to determine the basal level activities of ERα without any induction or drug treatment were done in hormone-deprived media. For the treatments with purine nucleosides, cells were incubated in a medium containing purine-depleted serum for 72 h and then treated with vehicle (DMSO), adenosine (5 μM), guanosine (5 μM), inosine (5 μM), or a combination of the three nucleosides.

#### B. RNA extraction and real-time RT-qPCR

RNA extraction was done using a homemade guanidinium-acid-phenol reagent (*101*). 0.5 x 10^6^ cells were lysed in 500 μl solution D (4 M guanidium isothiocyanate, 25 mM sodium citrate pH 7 and 0.5% N-laurosylsarcosine, and 0.1 M β-mercaptoethanol), and then transferred to microfuge tubes. 50 μl of 2 M sodium acetate pH 7.4, 500 μl water-saturated phenol (ROTH #A980.1), and 100 μl chloroform/isoamyl alcohol (49:1) (Merck, #25668) were added and tubes were shaken vigorously. After centrifugation (20 minutes at 10000 g, 4°C), the top aqueous phase (about 400 μl) containing the total RNA was recovered, precipitated with 250 μl isopropanol, and RNA pellets were washed with 250 μl of 75% ethanol. RNA was quantitated using a NanoDrop. 400 ng of the total RNA were reverse-transcribed into cDNA using random primers (Promega #C1181) and the GoScript reverse transcriptase kit (Promega #A5003). Quantitative real-time PCR (qPCR) reactions were done using specific primers and the GoTaq qPCR Master Mix kit (Promega #A6001), according to the manufacturer’s protocol. Ct values were measured using a Bio-Rad CFX 96 Real-Time PCR instrument (Bio-Rad). Relative gene expression was calculated using the ΔΔCt method. Specific primer sequences are listed in Table S2.

### Protein expression analyses

#### A. Drug treatments and transient transfections

Assays of the basal levels of ERα were done in a hormone-deprived medium to stabilize the receptor. For treatments with purine nucleosides, cells were pre-incubated in hormone-deprived medium for 72 h for ERα assays or in a medium containing purine-depleted serum for the assays of mTOR signaling markers. Then, cells were treated with vehicle (DMSO), adenosine, guanosine, inosine, or a combination of the three nucleosides. Nucleosides were added at 5 μM each for assays in purine-depleted medium, whereas excess nucleosides at 100 μM were used for assays in hormone-deprived but purine-rich medium. For CREB1 phosphorylation assays, cells were treated with FI cocktail that contains 10 μM forskolin (Cayman chemical #11018) and 100 μM IBMX (Sigma-Aldrich #410957), and incubated for 30 minutes, 4 h, 12 h, and 24 h. For assays with MRT, cells were treated/pre-treated with MRT (500 nM) for 24 h. For the transient expression of sgNT and sgPAICS in PAICS-overexpressing MCF7-V clones, cells were hormone-deprived for 72 h, seeded in a 6-well plate and transfected with either sgRPE65 or sgPAICS plasmids using PEI MAX 40K at a DNA:PEI MAX ratio of 1:6.

#### B. Immunoblot analyses

Cells were harvested and pellets were washed at least twice with PBS, and then lysed in ice-cold lysis buffer (20 mM Tris-HCl pH 7.4, 2 mM EDTA, 150 mM NaCl, 1.2% sodium deoxycholate, 1.2% Triton-X-100, protease inhibitor cocktail (Thermo Fisher Scientific #78429), and phosphatase inhibitor cocktail (Thermo Fisher Scientific #78420)). Cell lysates were sonicated for 15 min at high power with the Bioruptor sonicator (Diagenode). Protein quantification was performed using the Bradford reagent (Biorad #5000001), measuring absorbance at 595 nm. 20-50 μg of proteins were resolved on SDS-PAGE and transferred to a nitrocellulose membrane (GVS Life Science) with an electroblotting unit (VWR #BTV100). Membranes were blocked with 2.5% non-fat milk or BSA in TBS containing 0.2%Tween-20 (TBST), and incubated with primary antibodies overnight at 4°C. Then they were washed with TBST, incubated with the corresponding secondary antibodies for 1 hour at room temperature, and developed using the WesternBright^TM^ chemiluminescent substrate (Advansta #K-12045-D50). Images were captured using an Amersham™ ImageQuant™ 800 biomolecular imager. The “Fisher BioReagents™ EZ-Run™ Prestained Rec Protein Ladder” (Thermo Fisher Scientific) was used as protein molecular weight marker. Details of primary and secondary antibodies are listed in Table S3.

#### Puromycin labeling assay

WT MCF7-V cells, and the MCF7-V clones PAICS NC, PAICS OE 25, and PAICS OE 45 were seeded at 60% confluency in 10 cm dishes. MRT was added to the cells for 24 h followed by puromycin (1 μM) for specific periods. Cells were harvested and lysed in 20 mM Tris-HCl pH 7.4, 2 mM EDTA, 150 mM NaCl, 1.2% sodium deoxycholate, 1.2% Triton-X-100, and protease inhibitor cocktail.

Immunoblotting was done using an anti-puromycin antibody (Table S3).

#### Dual-luciferase reporter assays

For assays with HEK 293T cells, they were seeded at 60% confluency in 6-well plates. After hormone deprivation for 72 h, cells were co-transfected using PEI MAX 40K with 0.75 μg of the luciferase reporter plasmid ERE-Luc, 50 ng of Renilla luciferase expression plasmid pRL-CMV used as a transfection control, and 1 μg of either pPAICS-EGFP or pEGFP-C1 as a control plasmid. Each transfection mixture was further divided to receive 0.15 μg of either plasmids HEG0 (*95*) or its derivatives HEG0 S305A and HEG0 S104/106A, or their expression vector pSG5 (*96*) as the empty vector control. 24 h after transfection, cells were trypsinized and seeded into 96-well plates, then treated with either ethanol, or E2 (1 nM), or E2 + 4-OHT (100 nM), or E2 + ICI (100 nM) for another 24 h. For assays with MCF7-V cells, WT cells or the sgPAICS 38 knockdown clone were pre-incubated in a medium containing purine-depleted (Thermo Fisher Scientific #A3382001) or purine-supplemented (Thermo Fisher Scientific #A3160501) serum for 72 h. Then, cells were seeded in 6-well plates and co-transfected with 1 μg ERE-Luc and 50 ng pRL-CMV using PEI MAX 40K. After 24 h, cells were trypsinized, re-seeded into 96-well plates, and treated with vehicle (DMSO), adenosine (5 μM), guanosine (5 μM), inosine (5 μM), or a combination of the three nucleosides, in the presence and the absence of E2 (5 pM). For luciferase assays with CRE-Luc, WT MCF7-V cells, and the knockdown clones sgPAICS 28 and sgPAICS 38 were co-transfected with 1 μg CRE-Luc reporter plasmid and 50 ng pRL-CMV using PEI MAX 40K. After 24 h, cells were trypsinized, re-seeded into a 96-well plate, and treated with either FI or vehicle (DMSO) for another 24 h. Cells were lysed, and luciferase activities were measured with the dual-luciferase detection kit (Promega #E1910), using a bioluminescence plate reader (Cytation 3 Image Reader, Agilent). The activity of firefly luciferase was normalized to the activity of the Renilla luciferase.

#### ATP assay

ATP levels were measured with the CellTiter-Glo (CTG) luminescent assay (Promega #G7572) according to the manufacturer’s instructions. Briefly, cells were seeded at a density of 70,000 cells per well of 96-well plates. After 24 hours, CTG reagent was added, and the luminescence was measured with a Cytation 3 Image Reader from Agilent. The luminescence of control cells was set to 1.

#### cAMP-Glo assay

cAMP was measured using the cAMP-Glo™ assay kit (Promega #V1501) according to the manufacturer’s protocol. Briefly, cells were seeded at a density of 3,000 cells/well of a white opaque 384-well plate with a clear bottom (Corning #3765). After 24 h, media was replaced by serum-free medium containing FI and incubated for 30 minutes, 1 h, 3 h, and 6 h. Cells were lysed using the cAMP-Glo™ lysis buffer by shaking at room temperature for 15–30 minutes. cAMP detection solution containing the enzyme PKA was added, mixed, and incubated for 20 minutes at room temperature. Then, the Kinase-Glo® reagent was added, mixed, and incubated for 10 minutes at room temperature. Luminescence was measured using a Cytation 3 Image Reader from Agilent.

#### PKA-SPARK fluorescence imaging

Cells were seeded at 200,000 cells/well of a 6-well plate and transiently transfected with 1 μg pPKA-SPARK using the PEI MAX 40K transfection reagent. 24 h after transfection, cells were treated with FI for specified time points. Fluorescence imaging and image analysis was done on live cells using a Cytation 3 Image Reader (Agilent) the software Gen5.

#### 4-OHT and MRT dose-response curves

Cells were seeded into a 96-well plate 24 h before drug treatments. A serial dilution ranging from 10^−9^ M to 10^−4^ M for 4-OHT or MRT in DMEM medium without phenol red, containing 5% CHFBS, penicillin/streptomycin, L-glutamine, and E2 (5 pM) was prepared and added to the cells. For the combination of 4-OHT + MRT, 50 nM or 120 nM MRT were added to the serial dilutions of 4-OHT. After 7 days of 4-OHT and 4-OHT + MRT treatments, or 3 days of MRT monotherapy, cells were stained using crystal violet and absorbance was measured at 595 nm using a Cytation 3 Image Reader (Agilent). Dose-response curves were fitted by least squares regression with a variable slope (four parameters) using GraphPad Prism version 8.0.0. IC10, IC20, and IC50 values were obtained by interpolating the curves with a 95% confidence interval.

#### Kaplan–Meier analyses

Graphs for OS and RFS associated with *PAICS* mRNA expression in ERα+ (+/− tamoxifen treatment) and ERα-breast cancer datasets were generated using the Kaplan–Meier Plotter (*40*).

#### Statistical analyses

Data were analyzed using GraphPad Prism 8.0.0 and Microsoft Excel. Differences between two groups were analyzed with a two-tailed Students *t*-test. One-way ANOVA was used to analyze variations between tissue samples of the TMA scan. Two-way ANOVA was used to analyze differences of multiple variables between more than two groups. Error bars represent the standard deviation between replicates and *p*-values < 0.05 were considered statistically significant.

## Acknowledgments

We are grateful to the iGE3 genomics core facility of the University of Geneva for NGS services. We especially thank Mylène Docquier for helping with sequencing the sgRNA library and Dr. Nicolas Hulo for his help using the MAGeCK-VISPR platform. We thank Dimitri Moreau of the ACCESS Geneva core facility for his help with cell sorting using FACS. We thank the bioimaging core facility of the faculty of medicine at the University of Geneva for scanning the TMA slide. We especially thank Lilia Bernasconi for her help with immunostaining, François Prodon and Nicolas Liaudet for their help in image acquisition and data analysis. We thank Simon Osborne of LifeArc, UK, for providing the PAICS inhibitor MRT00252040. We thank Leonardo Scapozza and Kaushik Bhattacharya for their help and fruitful discussions. We thank Brian Rowan of the Tulane University School of Medicine, New Orleans, for providing the antibody for phospho-S104/106 of ERα. We are grateful to Eric Allémann and all other colleagues mentioned in the text for gifts of materials.

## Author contributions

D.H. and D.P. conceived the study; D.H. designed and performed most of the experiments, analyzed most of the data, prepared the figures and wrote the drafts; V.V. contributed to the secondary screen; D.P. supervised the work, contributed to the design of the experiments, wrote, and critically edited the manuscript. All authors have read and agreed to the published version of the manuscript.

## Data and materials availability

With one exception mentioned in the text (full data available on request), all data are available in the main text or the supplementary materials.

## Supplementary Materials

**Fig. S1.**
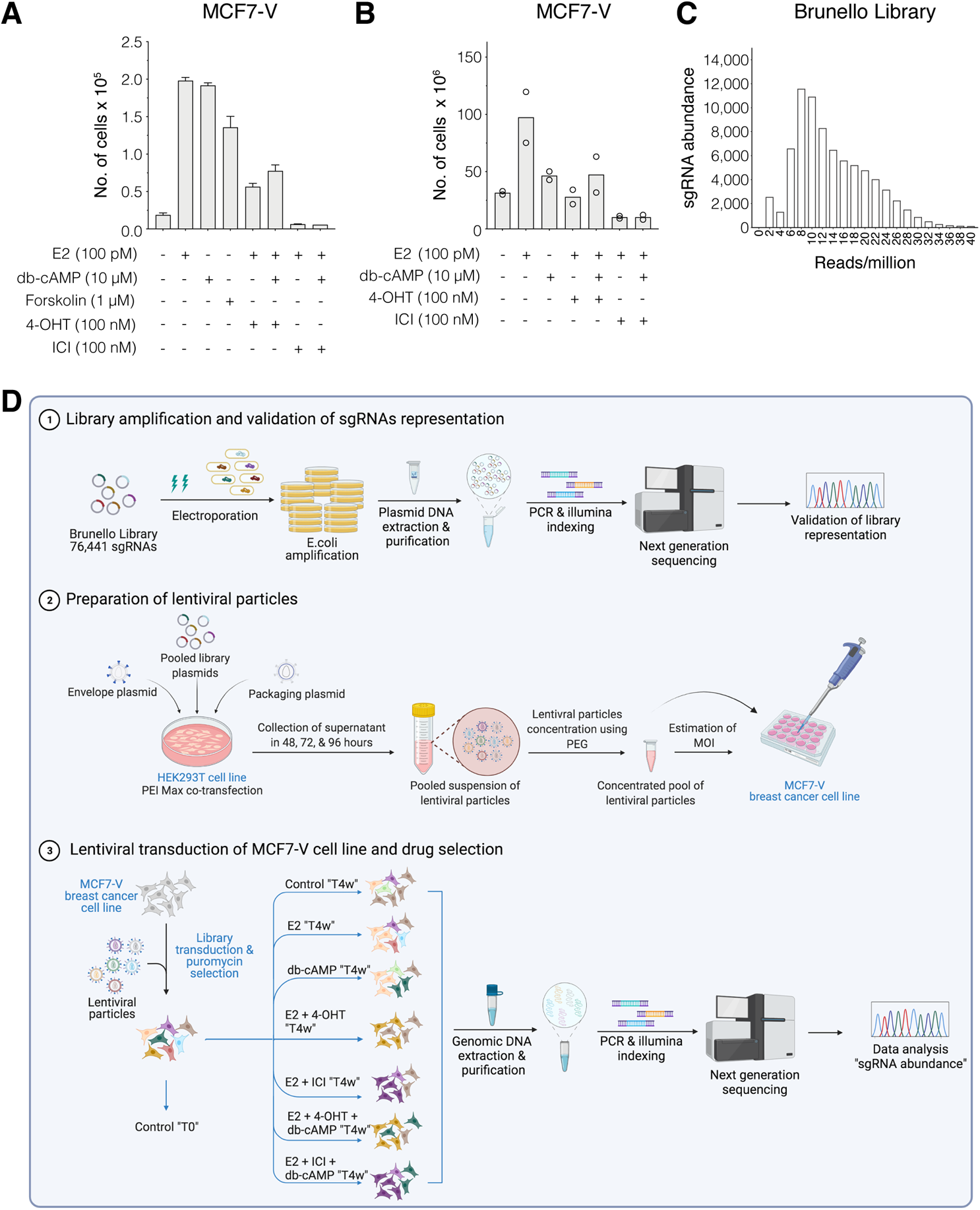
Details of CRISPR/Cas9 screen. **(A)** Bar graph representing the means of the number of MCF7-V cells after 2 weeks with the indicated treatments, measured by a crystal violet assay; cell numbers were interpolated from a standard curve. Error bars represent standard deviations of the means of at least 3 independent experiments. **(B)** Bar graph representing the cell counts of MCF7-V at the end of the primary CRISPR/Cas9 screen. Cells from two independent biological replicates were transduced with the Brunello library, treated as indicated for 4 weeks, and counted using a hemocytometer and an inverted light microscope. **(C)** Bar graph indicating the distribution and abundance of the sgRNAs after library amplification and NGS. **(D)** Schematic illustration of the detailed steps of the primary CRISPR/Cas9 screen.

**Fig. S2.**
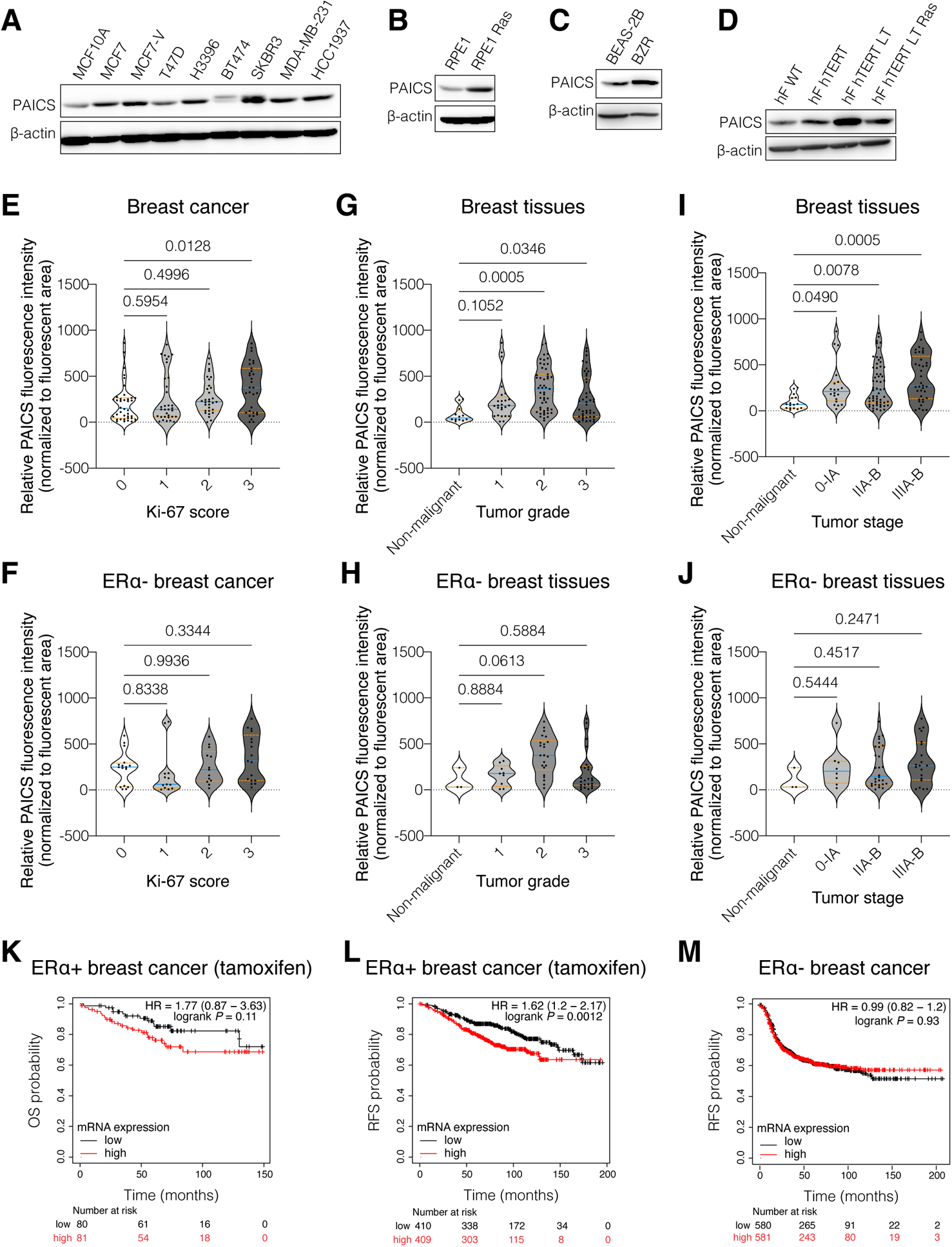
PAICS expression levels are upregulated in cancer, and correlate with progression and poor clinical outcomes of ERα+ breast tumors. **(A)** Immunoblot of PAICS in total cell lysates of a panel of breast cancer cell lines and the non-malignant cell line MCF10A. **(B to D)** Immunoblots of PAICS. **(E to J)** Violin plots of the means of the relative fluorescence intensities of PAICS per fluorescent unit area and its 95% confidence interval between cores classified based on Ki-67 score (panels E and F), tumor grade (panels G and H), and TNM stage (panels I and J). The statistical significance between the groups was analyzed by one-way ANOVA, and *p*-values < 0.05 were considered statistically significant. **(K** and **L)** Kaplan-Meier plots of the data of ERα+ breast cancer patients who received tamoxifen therapy, classified as tumors expressing high levels (red line) and low levels (black line) of *PAICS* mRNA. (**M)** Kaplan-Meier plots for the RFS of ERα-breast cancer patients.

**Fig. S3.**
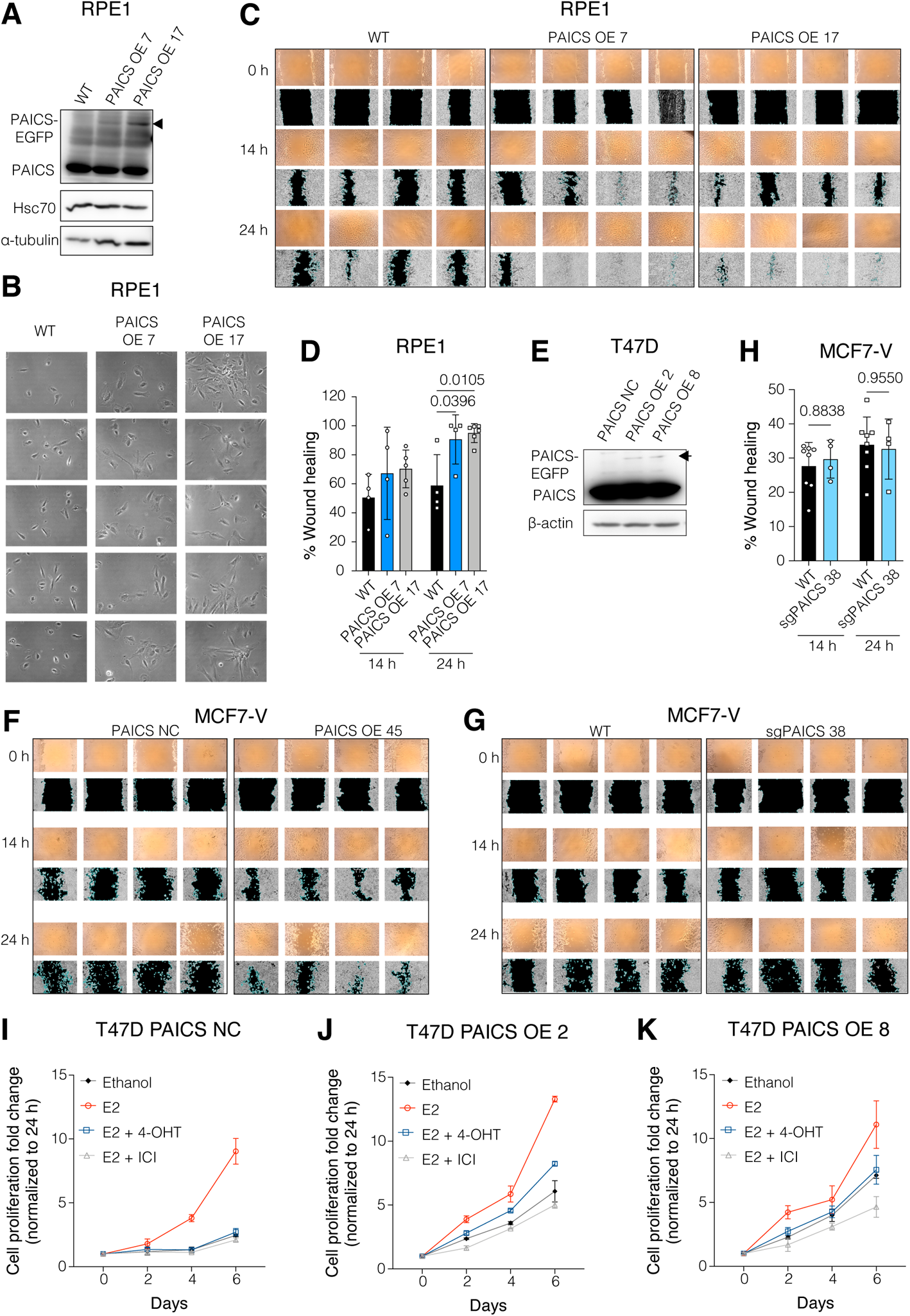
PAICS expression correlates with cell migration and estrogen-independent and tamoxifen-resistant proliferation of ERα+ breast cancer cells. **(A)** Immunoblots of endogenous PAICS and exogenous PAICS-EGFP. Hsc70 and α-tubulin were used as internal controls. **(B)** Cellular morphology of RPE1 cells as indicated. **(C, F** and **G)** Cell migration assays. Raw and ImageJ-processed images of the replicates of the scratch wound healing assays. Phase-contrast images were captured at the time of the scratch (0 h), after 14 h and 24 h using an inverted light microscope. **(D** and **H)** Bar graph showing the % wound healing as quantitated with the software ImageJ. **(E)** Immunoblots of endogenous PAICS and exogenous PAICS-EGFP expressed in T47D cell derivatives as indicated. **(I to K)** Cell proliferation of T47D cell derivatives as indicated, treated with either vehicle (ethanol), E2 (100 pM), E2 (100 pM) + 4-OHT (100 nM), or E2 (100 pM) + ICI (100 nM), measured with a crystal violet assay. For panels D, H, and I to K, data are represented as means ± SD of at least 3 independent experiments. The statistical significance between the groups was analyzed by two-way ANOVA (D and H), and *p*-values < 0.05 were considered statistically significant.

**Fig. S4.**
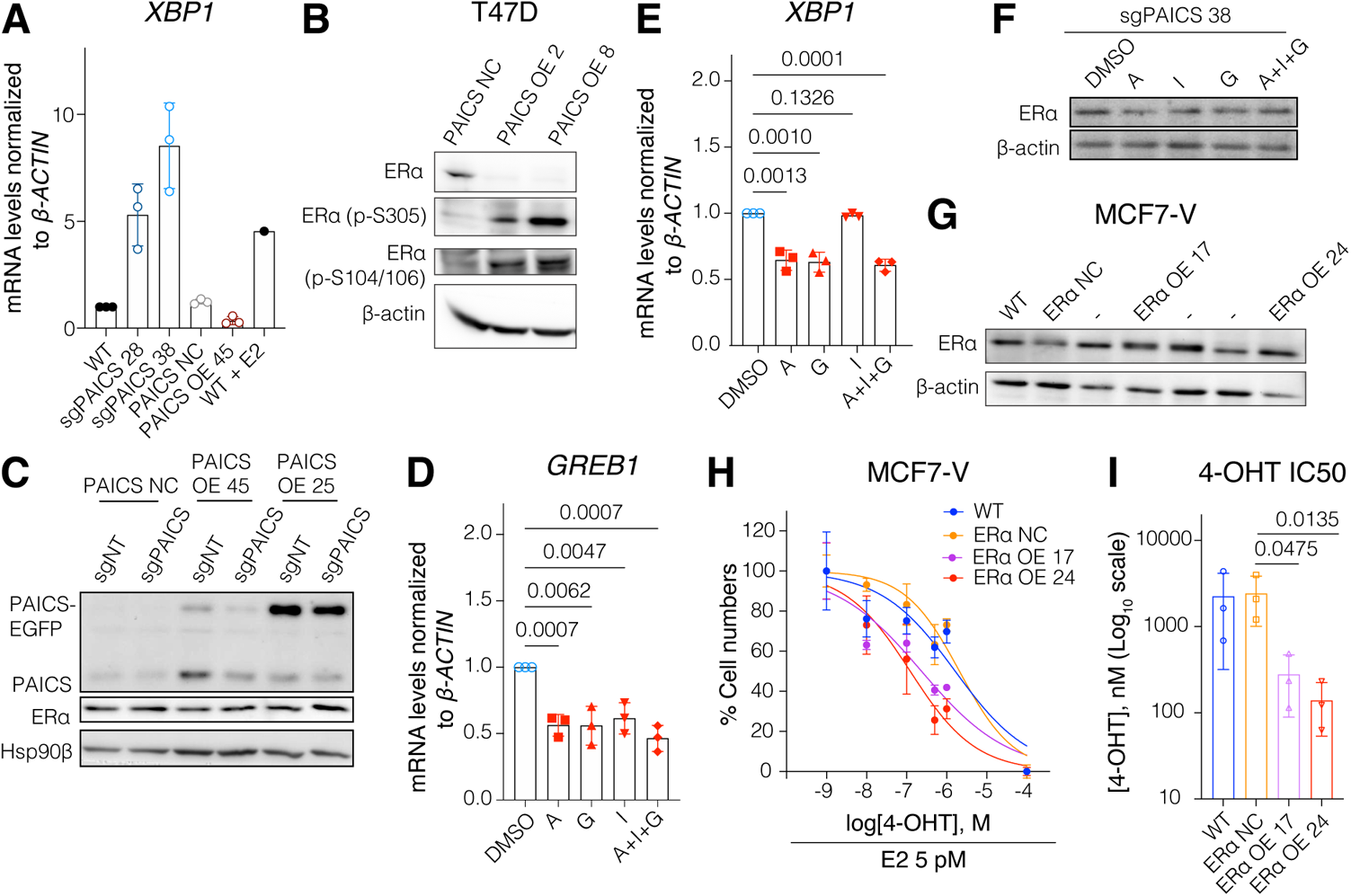
Interplay between PAICS and ERα. **(A)** mRNA levels of the ERα target gene *XBP1*. MCF7-V WT cells treated with E2 (1 nM) were considered as a positive control. Ct values were first normalized to *β-ACTIN*, then values corresponding to MCF7 WT were set to 1. **(B)** Immunoblots of phosphorylated (S305) and (S104/106), and total ERα in the indicated T47D cells (**C)** Immunoblots of PAICS and ERα. **(D** and **E)** mRNA levels of the ERα target genes *GREB1* and *XBP1* in MCF7-V cells. Cells were starved in medium containing purine-depleted serum for 72 h and then treated with either DMSO (vehicle), A (5 μM), G (5 μM), I (5 μM), or a combination of the 3 nucleosides (A + I + G) each at 5 μM. Ct values were first normalized to *β-ACTIN*, then values corresponding to vehicle were set to 1. **(F)** Immunoblot of ERα in the MCF7-V cell clone sgPAICS 38. Cells were hormone-starved for 72 h and then treated with either DMSO (vehicle), A (100 μM), G (100 μM), I (100 μM), or a combination of the 3 nucleosides (A + I + G) each at 100 μM. **(G)** Immunoblot of ERα. Before the assay, cells were hormone-starved for 72 h to stabilize ERα protein levels. **(H)** Dose-response curves with increasing doses of 4-OHT. **(I)** 4-OHT IC50 values interpolated from the 4-OHT dose-response curves shown in panel J; the data are represented as the interpolated means ± 95% CI. For panels C, D, F, G, and J, the data are represented as means ± SD of at least 3 independent experiments. The statistical significance between the groups was analyzed by unpaired Student’s t-tests (F, G, and K) or two-way ANOVA (C), and *p-*values < 0.05 were considered statistically significant.

**Fig. S5.**
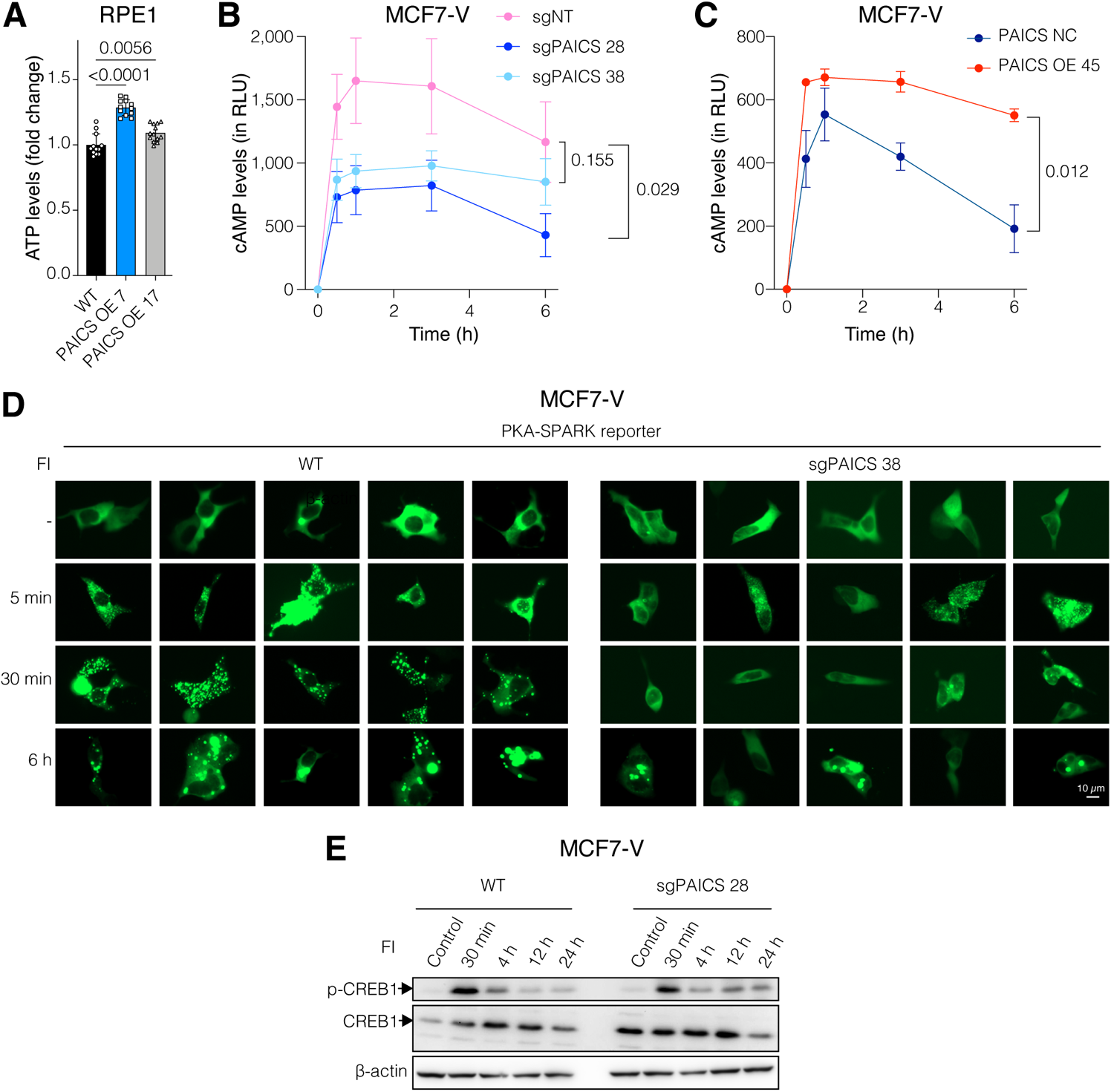
Increased expression of PAICS correlates with the activation of cAMP/PKA in ERα+ breast cancer cells. **(A)** ATP levels of indicated cells. Luminescence signal of RPE1 WT cells was set to 1. **(B** and **C)** cAMP levels measured by cAMP-Glo reagent. Cells were treated with FI for 30 min, 1 h, 3 h, and 6 h. Experiment was based on at least 3 independent replicates. **(D)** Fluorescent images of cells transiently transfected with the PKA-SPARK reporter, and treated with FI for 5 min, 30 min, and 6 h to induce cAMP/PKA signaling. Green-fluorescent phase separation droplets are indicative of active PKA. **(E)** Immunoblots of phosphorylated (S133) and total CREB1. Cells were treated with FI for 30 min, 4 h, 12 h, and 24 h to induce the phosphorylation of CREB1 downstream of active PKA. For panels A-C, data are represented as mean ± SD of at least 3 independent experiments. The statistical significance between the groups was analyzed by unpaired Student’s t-tests (A), and *p*-values < 0.05 were considered statistically significant.

**Fig. S6.**
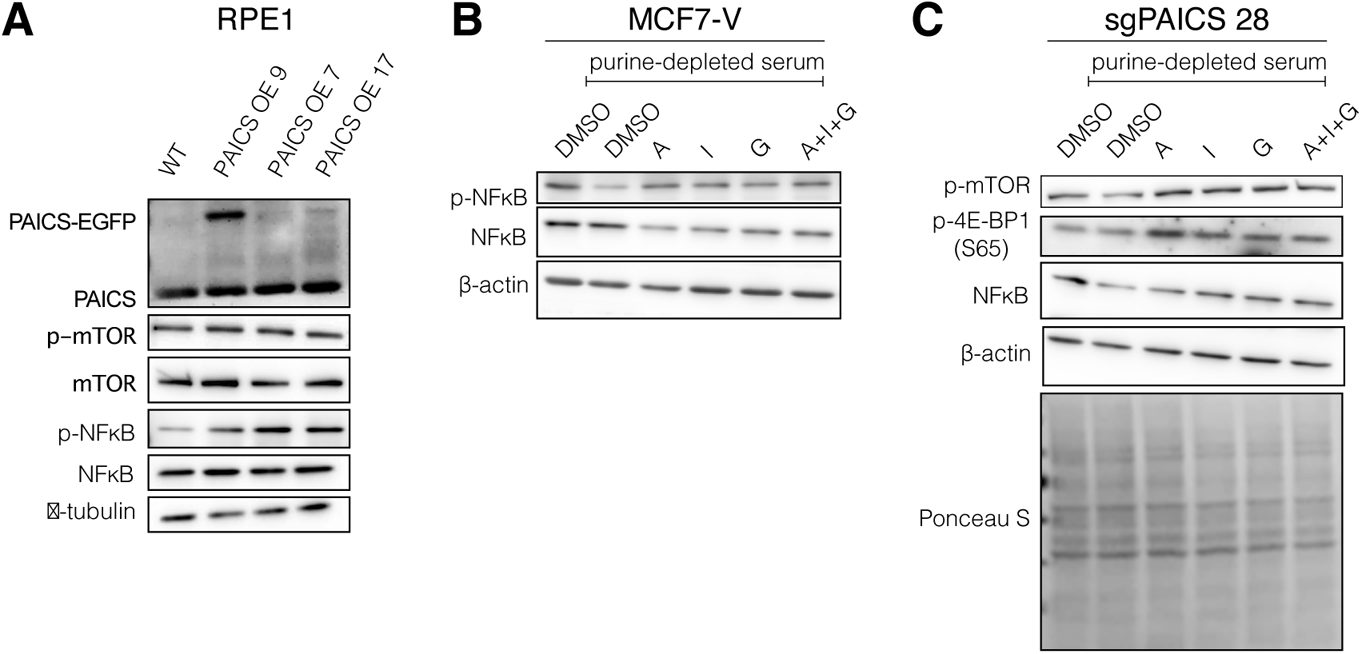
Expression of PAICS is correlated with mTOR activity and the global rate of protein translation. **(A)** Immunoblots of the indicated proteins. **(B** and **C)** Immunoblots of the indicated protein levels in MCF7-V WT (panels B) cells and in MCF7-V clone sgPAICS 28 (panel C). Cells were purine-depleted for 72 h and then treated with either DMSO (vehicle), A (5 μM), G (5 μM), I (5 μM), or a combination of the 3 nucleosides (A + I + G) each at 5 μM. In parallel, a control group of cells that were not purine-starved were incubated in purine-rich medium for 72 h and then treated with DMSO.

**Fig. S7.**
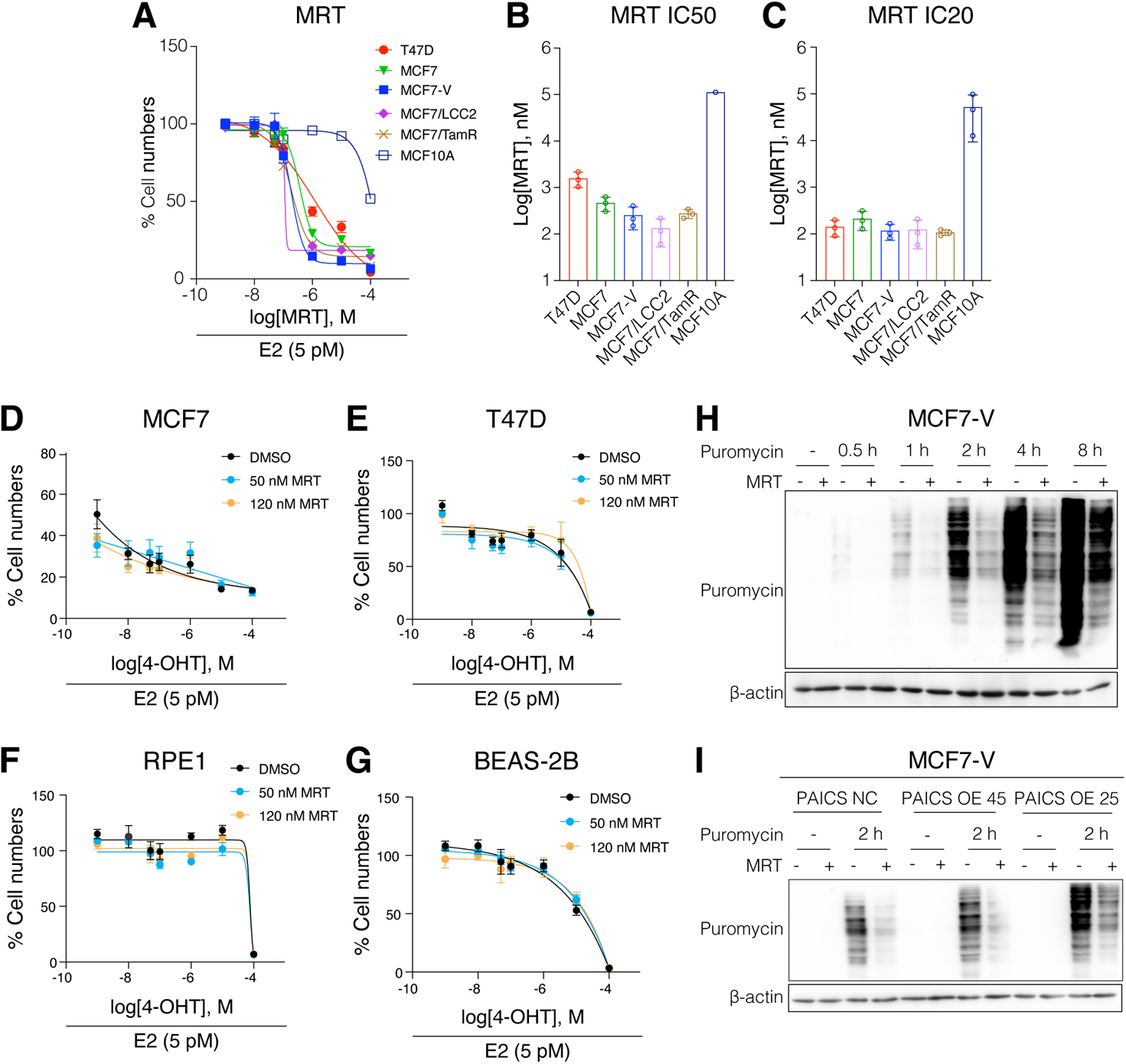
Pharmacological inhibition of PAICS sensitizes ERα+ breast cancer cells to 4-OHT. **(A)** Dose-response curves with increasing doses of MRT with the breast cancer cell lines T47D, MCF7, MCF7-V, MCF7/LCC2, and MCF7/TamR, and the non-malignant breast cells MCF10A. **(B** and **C)** Interpolated IC50 (panel B) and IC20 (panel C) values of MRT from the dose-response curves indicated in panel A. Data are represented as the interpolated means ± 95% CI. **(D to G)** Dose-response curves with increasing doses of 4-OHT with cell density measured with a crystal violet assay. For each curve, a control group treated with vehicle was set to 100%. **(H** and **I)** Immunoblot of puromycin-labelled nascent polypeptides. For panel H, MCF7-V WT cells were pre-treated with either vehicle (DMSO) or MRT (500 nM) for 24 h, then treated without or with puromycin (1 μM) for 0.5 h, 1 h, 2 h, 4 h, and 8 h. Cells in panel I were pre-treated with either vehicle (DMSO) or MRT (500 nM) for 24 h, then treated without or with puromycin (1 μM) for 2 h. For panels A, and D to G, data are represented as means ± SD of at least 3 independent experiments.

**Table S1.**
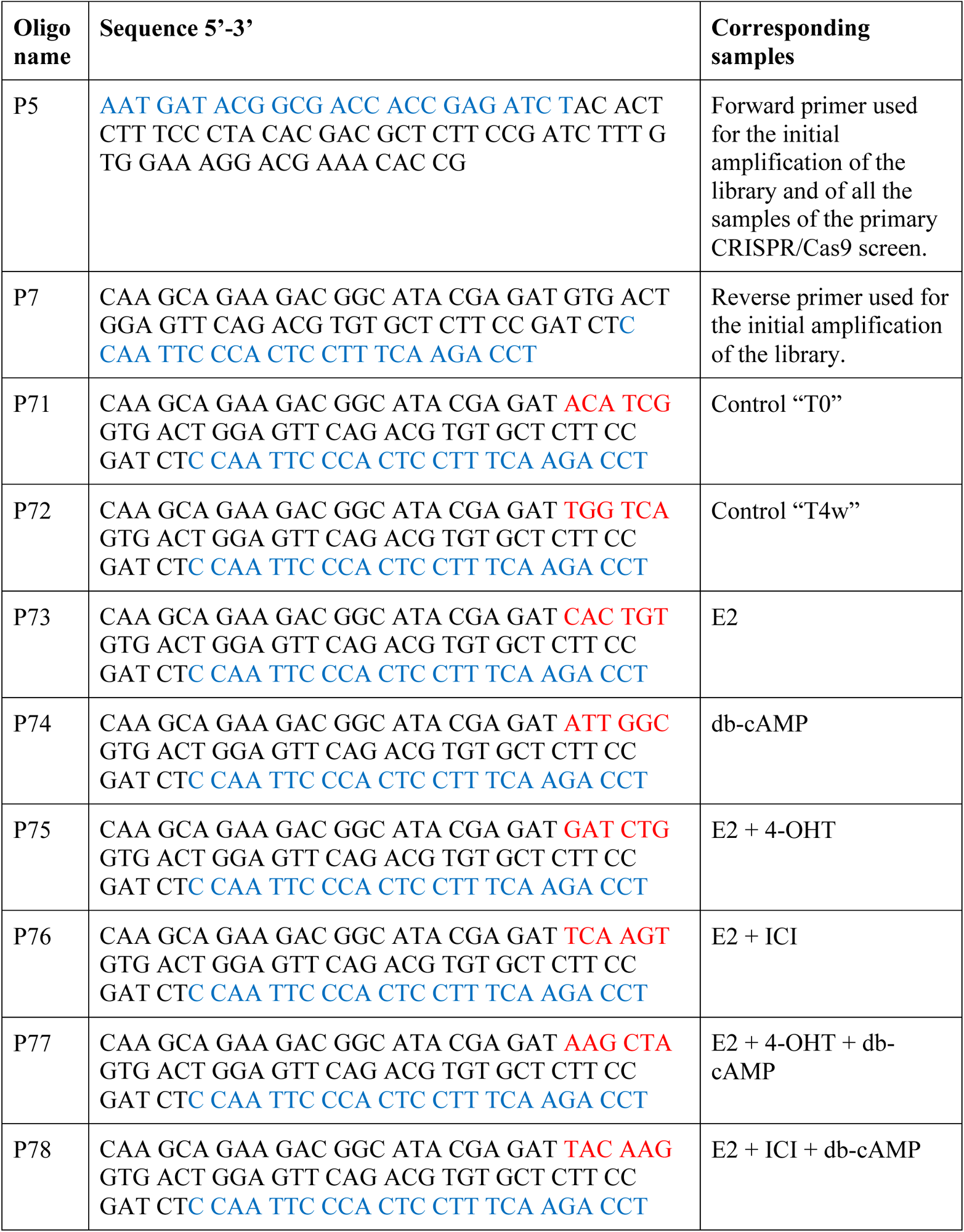
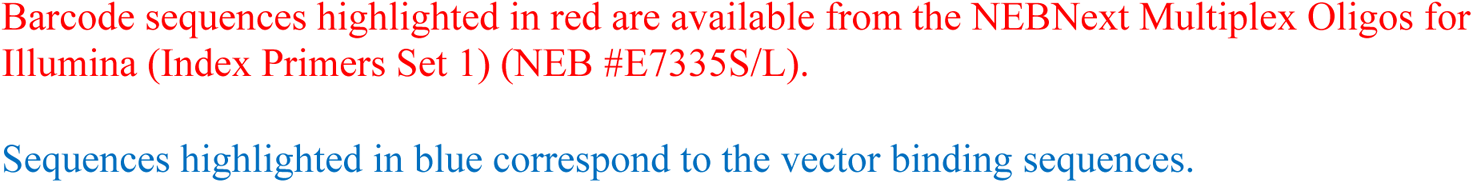
List of Illumina sequencing primers used for library amplification and NGS of the primary CRISPR/Cas9 screen.

**Table S2.**
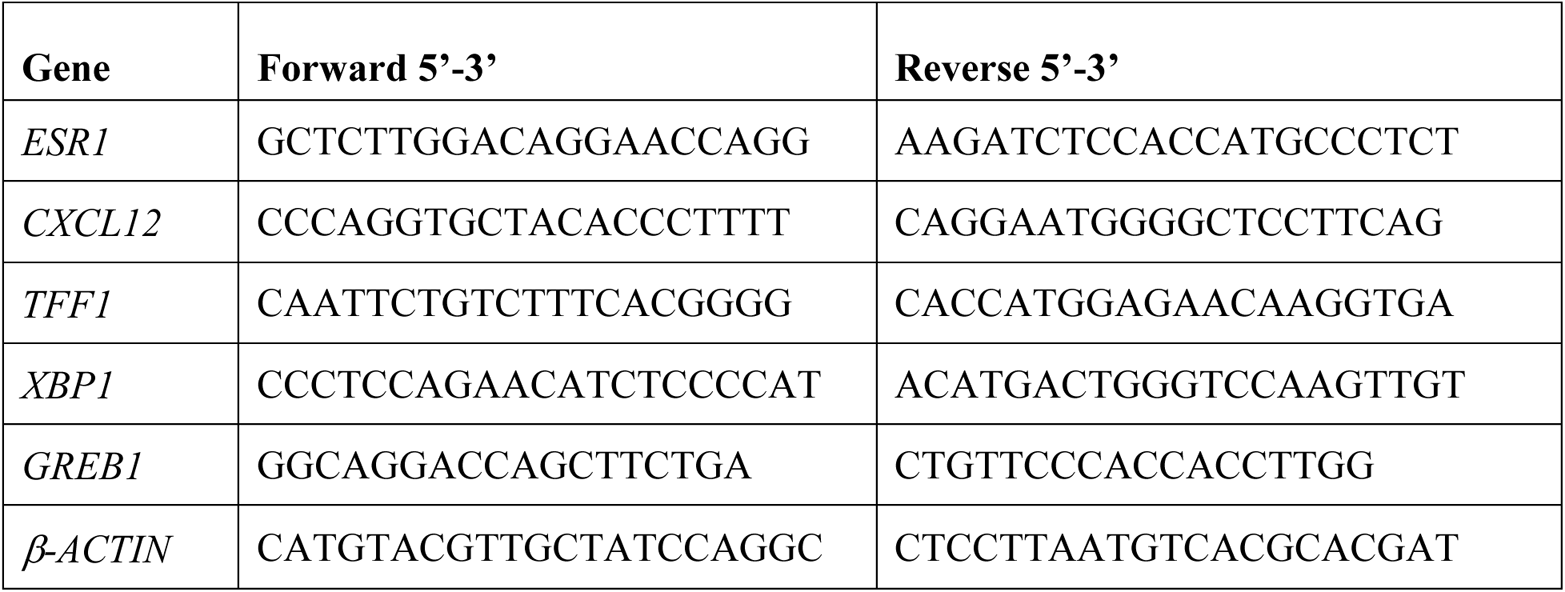
List of specific primer sequences used for real-time RT-qPCR.

**Table S3.** List of antibodies used in this study.

| **Antigen and/or antibody** | **Supplier** | **Catalog no.** | **Host** | **Clonality** | **Dilution** |
| --- | --- | --- | --- | --- | --- |
| ERα | Bethyl Laboratories | A300-498A-M | Rabbit | Polyclonal | 1:500 |
| PAICS | Atlas antibodies | HPA041538 | Rabbit | Polyclonal | 1:500 for western blotting |
|  |  |  |  |  | 1:25 for tissue staining |
| Phospho-S305 of ERα | Abnova | PAB12622 | Rabbit | Polyclonal | 1:300 |
| Phospho-S104/106 of ERα | Formerly available from Bethyl Laboratories) * | N/A | Rabbit | Polyclonal | 1:500 |
| CREB1 (48H2) | Cell signaling technology | 9197 | Rabbit | Monoclonal | 1:1000 |
| Phospho-S133 of CREB1 (87G3) | Cell signaling technology | 9198 | Rabbit | Monoclonal | 1:1000 |
| β-actin (13E5) | Cell signaling technology | 4970 | Rabbit | Monoclonal | 1:1000 |
| HSP73 against HSC70 | StressMarq | SMC-151 | Mouse | Monoclonal | 1:2000 |
| Hsp90β (scFv H90-10) | Geneva antibody facility | ABCD_AO870 | Mouse | Monoclonal scFV | 1:2000 |
| Puromycin (clone 12D10) | Sigma-Aldrich | MABE343 | Mouse | Monoclonal | 1:22000 |
| Phospho-S2448 of mTOR | Cell Signaling Technology | 2971 | Rabbit | Polyclonal | 1:1000 |
| mTOR (7C10) | Cell Signaling Technology | 2983 | Rabbit | Monoclonal | 1:2000 |
| RPS6 | GeneTex | GTX130450 | Rabbit | Polyclonal | 1:1000 |
| phospho- S235/236 of S6 ribosomal protein (D57.2.2E) XP® | Cell Signaling Technology | 4858 | Rabbit | Monoclonal | 1:1000 |
| NFκB p65 | GeneTex | GTX107678 | Rabbit | Polyclonal | 1:1000 |
| Phospho-S311 of NFκB p65 | Santa Cruz | sc-135769 | Mouse | Monoclonal | 1:1000 |
| 4EBP1 (N1C3) | GeneTex | GTX109162 | Rabbit | Polyclonal | 1:1000 |
| Phospho-S65 of 4E-BP1 | GeneTex | GTX133184 | Rabbit | Polyclonal | 1:1000 |
| phospho-S51 of eIF2α (119A11) | Cell Signaling Technology | 3597 | Rabbit | Monoclonal | 1:1000 |
| eIF2α | Cell Signaling Technology | 9722 | Rabbit | Polyclonal | 1:1000 |
| GFP from mouse IgG1κ (clones 7.1 and 13.1) | Roche | 11814460001 | Mouse | Monoclonal | 1:7500 |
| Mouse IgG (H+L), HRP secondary antibody | Invitrogen | 31430 | Goat | Polyclonal | 1:10000 |
| Rabbit IgG (H+L), HRP secondary antibody | Invitrogen | 31460 | Goat | Polyclonal | 1:10000 |
| Cy3 AffiniPure Anti-Rabbit IgG (H+L), secondary red fluorescent antibody | Jackson Laboratory | 711-165-152 | Donkey | Polyclonal | 1:1000 for tissue staining |

\* A gift from Brian Rowan (Tulane University School of Medicine, New Orleans).

**The following supplementary materials are available as separate files.**

**Data S1. Normalized counts of the sgRNAs of the primary CRISPR/Cas9 knockout screen.** Plain text document contains the raw read counts from the two biological replicates that were normalized per million reads and presented side by side.

**Data S2. Calculated β-score of sgRNAs of the primary CRISPR/Cas9 knockout screen.** Plain text document contains values that indicate the enrichment (β > 0) or depletion (β < 0) of sgRNAs in the corresponding treatment groups as compared to the control group at T0.

**Data S3. Genes essential for MCF7-V cell survival.** Microsoft excel workbook contains the list of 600 genes and the calculated β-scores of their corresponding sgRNAs. These genes were considered essential for survival since their sgRNAs were depleted regardless of the added treatments.

**Data S4. Top 58 differentially abundant sgRNAs between the treatment conditions.** CSV document contains the list of genes represented with the calculated β-score values of their corresponding sgRNAs. The 15 groups of the hierarchical clustering are indicated.

**Data S5. Oligonucleotide sequences used to construct the sgRNAs of the secondary CRISPR/Cas9 screen.** Microsoft excel workbook contains the sequences that were selected from the Brunello library based on the off-target score and the consistency of read counts between the two replicates of the primary screen.

**Data S6. Human breast tissue microarray.** Microsoft excel workbook contains the details of the tissue biopsies, including tissue type, pathology diagnosis, TNM stage, tumor grade, the status of ERα, PR, HER2, and Ki-67, the measured fluorescence intensities corresponding to PAICS, the respective fluorescent area, and the calculated relative fluorescence of PAICS to unit area.

